# ToRQuEMaDA: Tool for Retrieving Queried Eubacteria, Metadata and Dereplicating Assemblies

**DOI:** 10.1101/2020.11.15.363259

**Authors:** Raphaël R. Léonard, Marie Leleu, Mick Van Vlierberghe, Frédéric Kerff, Denis Baurain

## Abstract

TQMD is a tool which downloads, stores and produces lists of dereplicated prokaryotic genomes. It has been developed to counter the ever-growing number of prokaryotic genomes and their uneven taxonomic distribution. It is based on word-based alignment-free methods (*k*-mers), an iterative single-linkage approach and a divide-and-conquer strategy to remain both efficient and scalable. We studied the performance of TQMD by verifying the influence of its parameters and heuristics on the clustering outcome. We further compared TQMD to two other dereplication tools (dRep and Assembly-Dereplicator). Our results showed that TQMD is optimized to dereplicate at high taxonomic levels (phylum/class), whereas the other dereplication tools are optimized for lower taxonomic levels (species/strain), making TQMD complementary to the existing dereplicating tools. TQMD is available at <https://bitbucket.org/phylogeno/tqmd>.

## Introduction

The fast-growing number of available prokaryotic genomes, along with their uneven taxonomic distribution, is a problem when trying to assemble high-quality yet broadly sampled genome sets for phylogenomics and comparative genomics. Indeed, most of the new genomes belong to the same subset of hyper-sampled phyla, such as Proteobacteria and Firmicutes, or even to single species, such as *Escherichia coli* (6086 genomes as of November 2020), while the continuous flow of newly discovered phyla prompts for regular updates of in-house databases. This situation makes it difficult to maintain sets of representative genomes combining lesser known phyla, for which only few species are available, and sound subsets of highly abundant phyla. An automated straightforward method is required but would be far too slow if based on regular alignment algorithms.

Alignment-free methods are quantifiable ways of comparing the similarity of sequences without using an alignment ^1^. They have several advantages over alignment-based methods: they are computationally less expensive, they are resistant to gene shuffling and recombination events, and they do not depend on assumptions about sequence changes. In the review of Zielezinski et al. ^1^, two main categories of methods are described: the information theory-based methods and the word-based methods.

The rationale behind word-based methods is that similar sequences share a similar set of words. Sequence words are called *k*-mers and can be defined as all the words, of a given size *k*, that one can enumerate for a given alphabet. The idea is to compare the “dictionaries” of the words observed in two different genomes. If we compare a book and a copy of the same book, their dictionaries will be the same and thus considered redundant. That case would correspond to the comparison of strains from the same species with so few differences that they would be akin to typos in the book copy. If we compare the dictionaries of two books about the same subject, say two high fantasy novels with a similar story, their dictionaries will be similar but with more differences, as the setting differs. That case would correspond to differences between genomes of different genera or families. And if the subject is completely different, such as a novel in a high fantasy setting and a manual of macro-economy, their dictionaries will just have the basic language in common, with the specific vocabulary being completely different. That case would correspond to the difference between two phyla or even between a bacterium and an archaea.

The information theory-based methods compute the amount of information shared between two analyzed (genomic) sequences. Several different ways to assess this quantity do exist but we will only briefly discuss two of them. The Shannon entropy ^2–4^, where the idea is that some words are common and thus their presence is unsurprising but the presence of rarer words is meaningful. The uncertainty to find a word in the sequence (or a text) is computed then the “indexes” of two different sequences are compared. As in the example with the high fantasy novels, the more the “indexes” share the same meaningful words, the more they are related. The Shannon entropy can also be used to identify interesting parts in genes in order to focus on them in phylogenetic analyses. The entropy being an indication of the level of variability in the sequence, a high entropy is an indication of the presence of common words. By identifying then removing the gene regions with common words (high entropy), one can enhance the variability (and thus the phylogenetic signal), reducing in the same time the required computational time ^5^. Another metric is the complexity of a sequence, as defined by Kolmogorov ^6^, can be measured by the length of its shortest description. This measure is commonly approximated with compression algorithms. The idea is to concatenate the two sequences to be compared and then compress the concatenated sequence. If the two sequences are the same, the compressed size of the concatenation will be equal, or almost equal, to the size of a single compressed sequence. In contrast the more different the sequences are from each other, the more their combined compressed size will grow until they are so different that the compressed size of their concatenation is roughly equal to the sum of the compressed sizes of the two sequences compressed separately.

Based on the review on the alignment-free sequence comparison methods of Zielezinski et al. ^1^, two main categories of software packages were theoretically suitable for dereplicating prokaryotic genomes: the species identification/taxonomic profiling programs (Table 1 in ^1^) and the whole-genome phylogeny programs (Table 2 in ^1^). First, we did not investigate software solutions made available as web services because of their intrinsic limitation with respect to the amount of genomic data that one regular user can process through these interfaces. Second, all the programs belonging to the taxonomic profiling category required a reference database to compare the genomes to, which would have led us to a circular conundrum, in which a (possibly handmade) database of reference genomes is required to (automatically) build a database of representative genomes. Third, all those presented in the whole-genome phylogeny category were either not suited for large-scale dereplication or did not provide small enough running time estimates for their test cases. For example, jD2Stat ^7^ gives results for 5000 sequences of 1500 nucleotides in 14 minutes, which would clearly make computationally intractable the dereplication of 82,000 whole prokaryotic genomes.

**Table 1.** Details of TQMD runs and phylogenomic datasets built on eight different subsets of Bacteria. For each dataset, TQMD was launched with the Jaccard Index as a distance, a pack size of 200, and was allocated a maximum of 50 CPUs. Other parameters (direct or indirect approach and distance threshold) are provided in the table, along with the total running time in CPU hours (h.CPU), the initial number of genomes (# starting), the number of representatives obtained (# repr.), the number of ribosomal protein alignments used in the supermatrix (# prot.), and the number of unambiguously aligned amino acids in the supermatrix (# AA). Further details (taxonomy and download links, Krona taxonomic plots, Forty-Two reports, supermatrices and trees) are available at <https://figshare.com/s/b63ee5d8839b5a01463e>.

| label | dataset | approach | threshold | h.CPU | # starting | # repr. | # prot. | # AA |
| --- | --- | --- | --- | --- | --- | --- | --- | --- |
| A | Bacteria (49) | indirect | 0.100 | 656 | 63,863 | 49 | 53 | 6338 |
| B | Bacteria (151) | indirect | 0.120 | 656 | 63,863 | 151 | 53 | 6187 |
| C | Actinobacteria | direct | 0.100 | 96 | 8859 | 20 | 51 | 6562 |
| D | Bacteroidetes | direct | 0.150 | 16 | 1225 | 37 | 49 | 6605 |
| E | Chlamydia | direct | 0.200 | 6 | 360 | 32 | 44 | 6131 |
| F | Cyanobacteria | direct | 0.200 | 8 | 428 | 46 | 48 | 6314 |
| G | Firmicutes | direct | 0.100 | 242 | 21,544 | 22 | 52 | 6536 |
| H | Proteobacteria | direct | 0.115 | 310 | 30,690 | 36 | 53 | 6471 |

**Table 2.** Comparison of the clustering properties when varying the distance metric, the distance threshold or the clustering approach. Analyses were run on 63,863 RefSeq Bacteria using two different distance metrics, either the Jaccard Index (JI) or the Identical Genome Fraction (IGF), six different distance thresholds (from 0.1 to 0.2 and from 0.3 to 0.4, respectively), and two different clustering approaches, either direct (JI-D and IGF-D) or indirect (JI-i and IGF-i; see text for details). All pack sizes were 200. RI = Redundancy Index (# groups / # phyla).

| Jaccard Index (JI) |  |  |  |  |  |  |  |  |  |  |  |  |  |
| --- | --- | --- | --- | --- | --- | --- | --- | --- | --- | --- | --- | --- | --- |
| direct approach (JI-d) |  |  |  |  |  |  | indirect approach (JI-i) |  |  |  |  |  |  |
| threshold | 0.20 | 0.18 | 0.16 | 0.14 | 0.12 | 0.10 | threshold | 0.20 | 0.18 | 0.16 | 0.14 | 0.12 | 0.10 |
| RI | 59 | 47 | 35 | 25 | 14 | 10 | RI | 54 | 46 | 35 | 20 | 12 | 4 |
| # phyla | 34 | 34 | 29 | 24 | 19 | 11 | # phyla | 34 | 31 | 25 | 22 | 13 | 11 |
| # groups | 2005 | 1589 | 1025 | 598 | 268 | 109 | # groups | 1845 | 1430 | 870 | 446 | 151 | 49 |
| - pure groups | 1992 | 1576 | 1009 | 587 | 261 | 106 | - pure groups | 1835 | 1416 | 853 | 434 | 149 | 45 |
| -- singletons | 1201 | 904 | 557 | 325 | 143 | 56 | -- singletons | 1727 | 818 | 488 | 242 | 88 | 24 |
| - mixed groups | 13 | 13 | 16 | 11 | 7 | 3 | - mixed groups | 10 | 14 | 17 | 12 | 2 | 4 |
| -- paraphyletic | 0 | 0 | 0 | 0 | 0 | 0 | -- paraphyletic | 0 | 1 | 0 | 0 | 0 | 0 |
| -- super-phyla | 10 | 10 | 12 | 5 | 2 | 0 | -- super-phyla | 10 | 13 | 9 | 7 | 0 | 1 |
| -- polyphyletic | 3 | 3 | 4 | 6 | 5 | 3 | -- polyphyletic | 0 | 0 | 8 | 5 | 2 | 3 |
| Identical Genome Fraction (IGF) |  |  |  |  |  |  |  |  |  |  |  |  |  |
| direct approach (IGF-d) |  |  |  |  |  |  | indirect approach (IGF-i) |  |  |  |  |  |  |
| threshold | 0.40 | 0.38 | 0.36 | 0.34 | 0.32 | 0.30 | threshold | 0.40 | 0.38 | 0.36 | 0.34 | 0.32 | 0.30 |
| RI | 74 | 65 | 58 | 45 | 33 | 31 | RI | 50 | 55 | 44 | 30 | 23 | 11 |
| # phyla | 24 | 24 | 22 | 22 | 22 | 15 | # phyla | 31 | 25 | 24 | 24 | 19 | 16 |
| # groups | 1776 | 1548 | 1271 | 988 | 719 | 464 | # groups | 1536 | 1369 | 1061 | 715 | 440 | 176 |
| - pure groups | 1758 | 1530 | 1271 | 971 | 706 | 456 | - pure groups | 1514 | 1345 | 1042 | 701 | 426 | 167 |
| -- singletons | 1094 | 939 | 755 | 587 | 419 | 260 | -- singletons | 905 | 784 | 595 | 404 | 219 | 77 |
| - mixed groups | 18 | 18 | 19 | 17 | 13 | 8 | - mixed groups | 22 | 24 | 19 | 14 | 14 | 9 |
| -- paraphyletic | 4 | 2 | 2 | 2 | 1 | 1 | -- paraphyletic | 2 | 3 | 1 | 0 | 2 | 0 |
| -- super-phyla | 11 | 11 | 13 | 10 | 4 | 1 | -- super-phyla | 17 | 17 | 14 | 10 | 8 | 4 |
| -- polyphyletic | 3 | 5 | 4 | 5 | 8 | 6 | -- polyphyletic | 3 | 4 | 4 | 4 | 4 | 5 |

Considering the limitations of the existing tools for assembling representative sets of prokaryotic genomes, the present article describes our own program called “ToRQuEMaDA” (abbreviated TQMD) for Tool for Retrieving Queried Eubacteria, Metadata and Dereplicating Assemblies. TQMD is a word-based alignment-free dereplicating tool for prokaryotic genomes designed for high-performance computing (HPC) environments. TQMD is available on BitBucket and can be installed on any HPC with SGE/OGE (Sun/Open Grid Engine) installed as a scheduler. Few modifications are needed to adapt the scripts to most local setups. TQMD works both in parallel and iteratively. Using default parameter values, each elemental job takes two to three hours to complete (see Materials and Methods for test hardware specifications), and if enough CPUs are available to run all jobs of a given round at the same time, such a round should only take two to three hours. Usually, four to five rounds are sufficient to achieve the dereplication. Therefore, a single run of TQMD against all Bacteria in NCBI RefSeq takes 8 to 15 hours to complete.

## Materials and Methods

### Hardware

Almost all the computational work was performed on a grid computer IBM/Lenovo Flex System composed of one big computing node (x440) and nine smaller computing nodes (x240), featuring a total of 196 physical cores, 2.5 TB of RAM and 160 TB of shared mass storage, and operating under CentOS 6.6. Beyond “bignode” (running the scheduler and the MySQL database; see below), only four of the smaller computing nodes were used when testing TQMD; their specifications are as follow: 2 CPUs Intel Xeon E5-2670 (8 cores at 2.6 GHz with Hyper-Threading enabled), 128 GB of RAM. For the dRep test (see below), we had to use a desktop workstation (Ubuntu Linux 16.04) featuring 2 CPUs Intel Xeon E5-2620 v4 (8 cores at 2.1 GHz with Hyper-Threading enabled) and 64 GB of RAM. Based on the comparator found on the website http://cpubenchmark.net/, the CPUs in our cluster and in the workstation were roughly equivalent (from −0.5 to +5% difference).

It is important to mention that due to Hyper-Threading configuration of the grid computer and the fact that several teams shared the infrastructure, queueing time and disk usage could not be strictly controlled during the tests. Therefore, all running times provided in this article are informed estimates rather than exact measures. These estimates are those we would communicate to a user inquiring about the waiting time for a specific analysis to complete. They are an approximation of the running time recorded when the grid computer usage is low (i.e., almost no other user).

### Software architecture

TQMD is composed of a database and includes two main phases: (1) a periodic preparation phase in which newly available genome assemblies (“genomes” for short) are downloaded and individual genome metrics are computed, and (2) an “on-demand” dereplicating phase in which genomes (both new and old) are dereplicated on the fly to provide a list of high-quality representative genomes as a result [Figure 1]. The database stores the paths to the individual genome (FASTA) files, the individual genome metrics and the list of representative genomes produced by each TQMD run. Each piece of data is computed independently; if a dereplication request is issued during the computation of newly available genomes, TQMD only uses the genomes for which all the data is available in the database.

**Figure 1.**
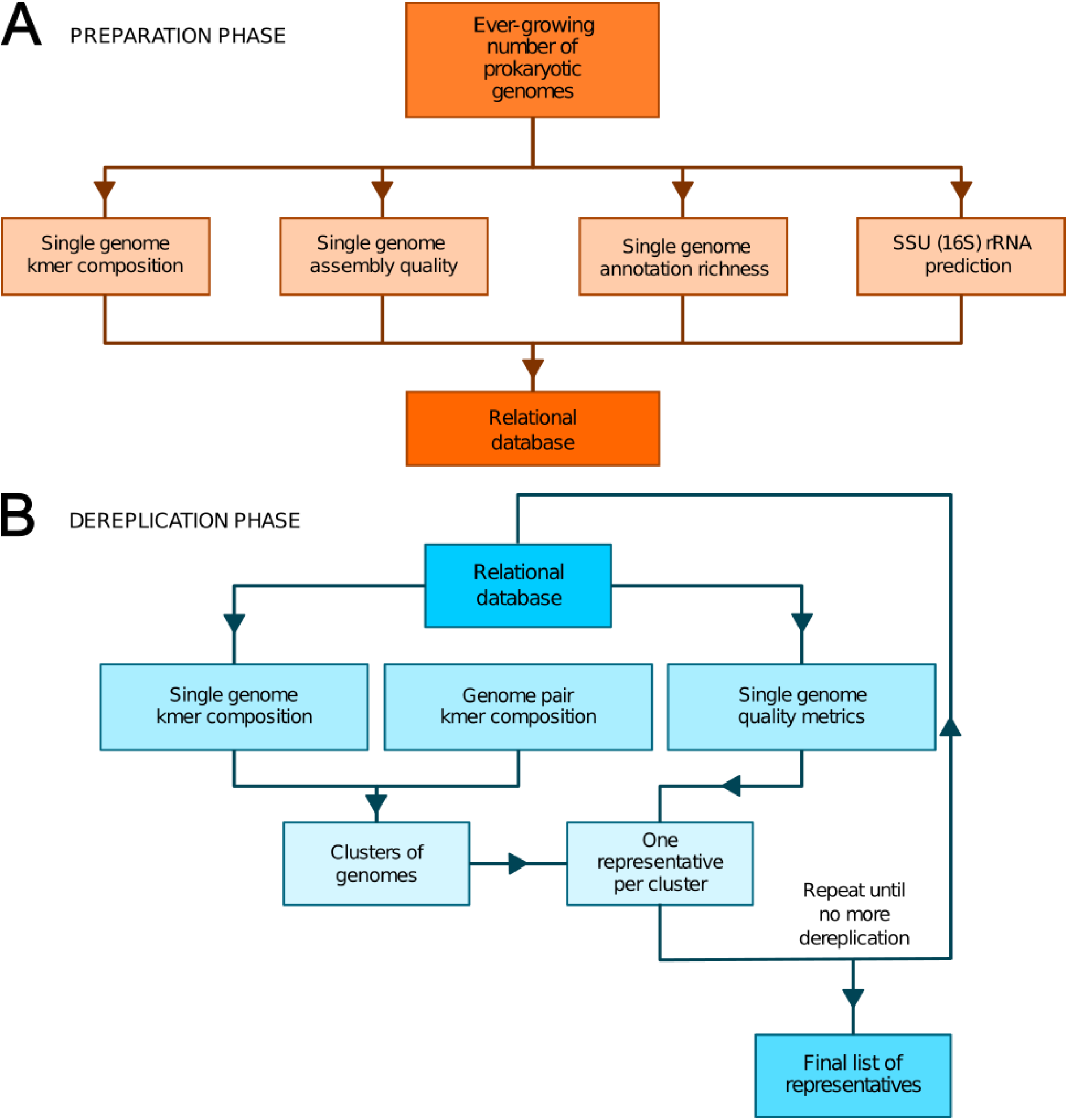
Overview of TQMD phases and heuristics. **A.** The preparation phase consists in downloading newly released prokaryotic genomes from NCBI RefSeq and to pre-compute all per-genome information required to run the second phase (k-mer composition, assembly quality, annotation richness and SSU rRNA (16S) prediction). Pre-computed information for single genomes is stored in a relational database associated with TQMD. **B.** The dereplication phase then retrieves this information for all genomes to dereplicate from the database and clusters the genomes from pairwise distances computed on the fly. Cluster representatives (one per cluster created) are chosen for each cluster based on the single genome metrics computed during the preparation phase. The dereplication is iterative and the process repeats until representative genomes cannot be dereplicated anymore, which produces the final list of representatives.

During the preparation phase, we pre-compute all the data required by the dereplication phase to store them in the database: indexes of nucleotide *k*-mers for single genomes, genome assembly quality metrics, genome annotation richness metrics and Small SubUnit ribosomal RNA (SSU rRNA 16S) predictions, whereas during the dereplication phase, we cluster the genomes based on these *k*-mer indexes and select a representative for each cluster based on assembly quality, annotation richness, and SSU rRNA 16S predictions. These criteria were chosen, so as to select the best representative genomes ^8^. By that, we mean that representative genomes (if available) are expected to be fully sequenced, correctly assembled, richly annotated and devoid of contaminating sequences. To satisfy this last requirement, TQMD can also use optional contamination statistics produced by Forty-Two ^9,10^ (see below).

#### Preparation phase

As shown in Figure 1A, we first download the prokaryotic genomes from NCBI RefSeq ^11^. For the sake of data traceability, TQMD never gets rid of older genomes; newly released genomes are simply added to its internal database. As TQMD was developed over 5 years, we have progressively accumulated several different versions of the RefSeq database, starting with release 79 (85,465 prokaryotic genomes, including 713 Archaea), then 79+92 (126,959 prokaryotic genomes, including 1037 Archaea) and finally 79+92+202 (155,519 prokaryotic genomes, including 1304 Archaea). Once RefSeq is up to date locally, we compute single-genome *k*-mer indexes and other metrics. For each of these computations, we use third-party programs and scripts (JELLYFISH, QUAST, RNAmmer, CD-HIT and Forty-Two), except for the richness of the annotations, which we evaluate using in-house scripts.

JELLYFISH (v1.1.12) ^12^ is used to compute the *k*-mer indexes for single genomes (TQMD can also work with JELLYFISH v2.x). We tested several sizes for our *k*-mers. However, we noticed that JELLYFISH v1.x crashes when using a size below 11 nucleotides, thus setting a hard lower bound on *k*-mer size. On the other hand, while longer k-mers improve the specificity, they also require longer computing times ^1^. With a size of 11, there are almost 4.2 millions (4^11^) possible words. Consequently, a hypothetical genome featuring every possible *k*-mer without any repetition, could only be 4.2 Mbp long. Even if real genomes include repeats, genomes over 4 Mbp might still feature almost every *k*-mer, which would lead to useless *k*-mer indexes. To verify this idea, we examined the 85,465 genomes of RefSeq 79 and observed that about 15 genomes indeed almost exhaust the *k*-mer index (3 to 4 millions out of 4.2 millions), thus confirming that 11 is not a usable *k*-mer size. The next *k*-mer size, 12 nucleotides, offers over 16 millions (4^12) possibilities. The genomes with the largest index only reach 7.5 millions different *k*-mers, while the average index is below 2.7 millions *k*-mers. We could have used a *k*-mer size of 13 nucleotides, but our preliminary tests showed an important increase of the computing time. Whereas our tests with a *k*-mer size of 12 on all available Bacteria lasted between 8 and 15 hours, depending on the distance threshold used (see below), our tests with a size of 13 required between 1 and 2 days to finish. Therefore, we chose to work with a *k*-mer size of 12 nucleotides. Above that, we would only have dereplicated closely related strains (i.e., belonging to the same pecies) due to a too high specificity ^1^ and/or the computing times would have become prohibitively long.

QUAST (v4.4) ^13^ is used to estimate the quality of genome assemblies. We retrieve several quality metrics (13 in total) for each genome, these being the number of DNA sequences, the number of DNA sequences (or contigs) > 1 kbp, the size of the complete genome, the size of the complete genome composed of DNA sequences > 1 kbp, the number of contigs, the largest DNA sequence, the size of the complete genome composed of DNA sequences > 500 bp, the GC content, the N50, N75, L50 and L75 values, and the number of “N” per 100 kbp (N is the symbol used for scaffold contigs without matching ends). Given a minimal set of contigs ordered by descending length, the N50/N75 is defined as the length of the contig located at 50%/75% of the total genome length in the distribution, whereas the L50/L75 is defined as the rank of this specific contig. Among these metrics, we eventually decided to take into account (1) the relative length of the largest DNA sequence to the complete genome (> 1 kpb only) and (2) the fraction of “N” in the genome. In addition, we also use a size range (between 100 kbp and 15 Mbp) to remove the genomes too small to be complete and those too large to be considered uncontaminated ^14^.

For the richness of annotation, we compute what we call the “certainty” and the “completeness” of each genome. Importantly, this step necessitates (predicted) proteomes. While it is not an issue with RefSeq genomes, for which such predictions are always available, if TQMD is provided with a dataset from a different source with missing predicted proteomes, the related genomes will be automatically discarded. Our “certainty” metric corresponds to the proportion of sequences in a given proteome that we deem uncertain. To this end, we first count the number of sequence descriptions (in FASTA definition lines) with words indicating uncertainty, such as “probable”, “hypothetical” or “unknown”, then we compute a relative score as follows:

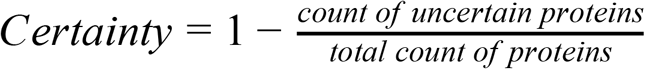

For “completeness”, instead of counting the number of uncertain proteins, we count the number of proteins without any description:

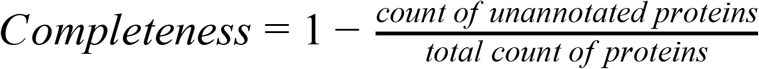

Regarding genome contamination, RNAmmer (v1.2) ^15^ is used to predict the SSU rRNA 16S of the genomes. By using cd-hit-est (v4.6) ^16,17^ with an identity threshold of 97.5% ^18^, we create a list of genomes featuring at least two SSU rRNA 16S sequences belonging to different species (i.e., clustered in distinct CD-HIT clusters). Note that another (more recent) possible threshold for species delineation based on SSU rRNA 16S identity would be 99% ^19^. In any case, this list of likely chimerical (or at least contaminated) genomes can be provided to filter out the genomes given as input to TQMD or produced in output by TQMD. Moreover, the presence of at least one SSU rRNA 16S predicted by RNAmmer is a requirement for the genome to be selected, which could rule out some metagenomes, for which rRNA genes are often missing ^20^.

Finally, another contamination metric is also available for the ranking but is not yet part of the automatic preparation pipeline of TQMD: the genome contamination level estimated by the program Forty-Two (v0.190820 or higher “42”) based on the comparison of the genome ribosomal proteins to the reference sequences of the RiboDB database ^21^.

Once all these individual *k*-mer indexes and metrics are computed for all individual genomes, the genomes are ranked in a global ranking from the best to the worst genome (to be selected as a cluster representative), using an equal-weight sum-of-ranks approach available in the Perl module Statistics::Data::Rank. For now, we do not use all the metrics available in the TQMD database. The five metrics that are actually used to compute the ranking are: (1) assembly quality: 100000 - number.Ns.per.100kbp; (2) assembly quality: largest.contig / total.length.in.1kbp.contigs; (3) annotation richness: certainty; (4) contamination level :100 - global.contamination.percentage; (5) number.enriched.alis / 90. The first two metrics are obtained from QUAST, the third from our in-house script and the two last from “42”, the latter two being optional.

#### Dereplication phase

As shown in Figure 1B, the dereplication process is iterative and stops once it deems itself finished. Its decision is based on three different convergence criteria, for which we provide default threshold values but these can be modified by the user (see below). TQMD stops cycling as soon as one criterion is satisfied.

Two different distances can be used for clustering genomes: the Jaccard index (JI; see ^22^) or the Identical Genome Fraction (IGF; see ^14^), both applied to shared *k*-mers at the nucleotide level.

The Jaccard index is a measure of the similarity between two finite datasets. It is defined as the intersection over the union of the two datasets A and B:

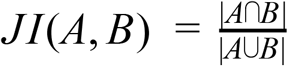

The IGF, for Identical Genome Fraction, replaces the union in the Jaccard index by the size of the smallest of the two datasets A and B:

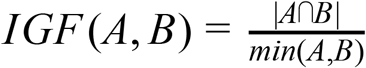

The TQMD algorithm works similarly for both distances and is inspired by the greedy clustering approach implemented in packages such as CD-HIT ^16,17,23^. First, we sort the list of genomes based on the global ranking of the genomes (assembly quality and annotation richness metrics, indicators of genome contamination; see above for details) and the top-ranking genome is assigned to a first cluster. Then every other genome is compared to every member of every cluster until it finds a suitable cluster of similar genomes; otherwise, such a genome becomes the first member of a new cluster. Hence, the second genome is compared to the single genome of the first cluster. If its distance to the latter genome is lower than specified threshold (let us say it is the case here), it is added to the cluster. Similarly, the third genome is compared to the first member of the first cluster; if its distance is higher than the threshold, it is compared to the second member of this first cluster. If it still is higher than the threshold, and since there is no other cluster, it creates a new (second) cluster. The fourth genome follows the same path, as will all remaining genomes do until every genome of the list is assigned to a cluster, whether singleton or part of a larger group. As genomes are processed from the best to the worst in the global ranking, representative genomes (which correspond to cluster founders) are automatically the best possible for each cluster.

To scale up the greedy clustering algorithm, we used a divide-and-conquer approach ^23,24^ (Figure 2). Indeed, when performing our own tests, we worked with about 112,000 genomes, a number making clearly impossible to compare all genomes at once. Therefore, we first partitioned the list of genomes into smaller batches (hereafter termed “packs”) of 200 by default, either based on their advertised (NCBI) taxonomy ^25,26^ or completely at random. The clustering of each small pack yields a single representative, which we regroup into a new (shorter) list of genomes that is processed iteratively following the same algorithm. In the next round, only the selected representatives are compared between each other, thereby precluding the genomes that were not selected to be directly compared. While this heuristic results in an important speed-up, it may also prevent similar genomes to be mutually dereplicated because they were processed in distinct packs and replaced by representatives that are less similar. The iterative algorithm stops based on any of the following three criteria (which can be specified by the user): (1) if it reaches a maximum number of rounds, (2) if it falls below an upper limit for the number of representatives (i.e., number of clusters) or (3) if the clustering ratio between two successive rounds falls below a minimum threshold. We define the clustering ratio as the percentage of genomes dereplicated at the end of a TQMD round compared to the number of genomes still in the game at the beginning of the round.

**Figure 2.**
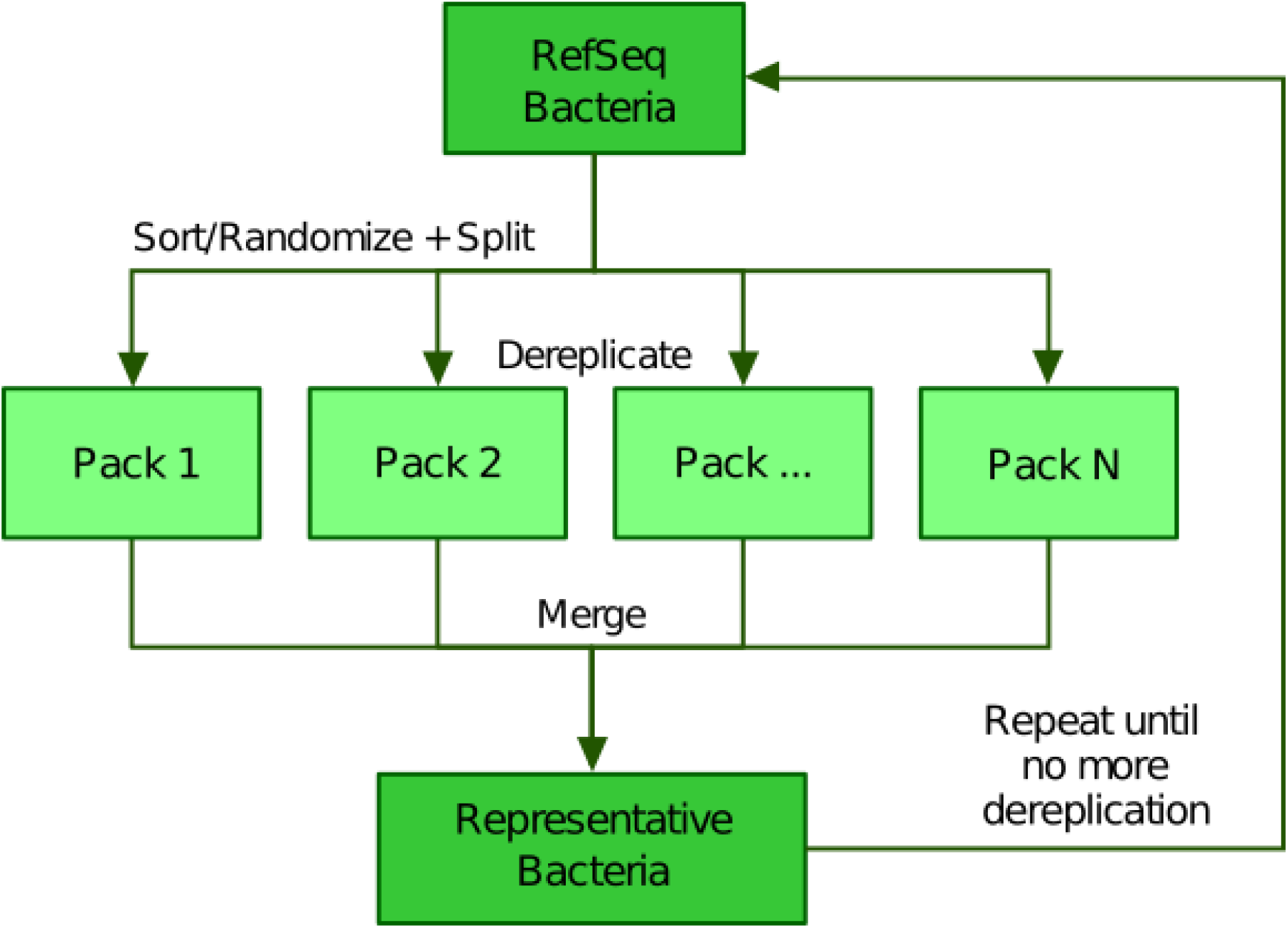
Illustration of the divide and conquer strategy of the dereplication phase. From a list of Bacteria downloaded from RefSeq, TQMD either sorts (based on the taxonomic lineage of each genome) or randomizes the list and splits it into packs of a given size. This allows each pack to be separately dereplicated, especially in parallel. Then all resulting lists of representative genomes are merged back and TQMD decides if it can stop or must refeed the merged list for another round.

### Phylogenomic analyses

We used TQMD runs as a source of representative bacterial genomes and obtained selections containing between 20 and 50 organisms for the six most populated phylums (the upper limit for the number of representatives was set to 50). We also generated two other selections to sample all Bacteria at once, one containing 49 organisms and the other 151. For each TQMD run, we retrieved the proteomes of the selected representatives and used Forty-Two to retrieve their ribosomal proteins. Those proteins were taxonomically labelled by computing the last common ancestor of their closest relatives (best BLAST hits) in the corresponding alignments (excluding self-matches), provided they had a bit-score ≥80 and were within 99% of the bit-score of the first hit (MEGAN-like algorithm ^14^). Thus, this strategy allowed us to simultaneously assess the completeness and the contamination level of each representative proteome while providing widely sampled ribosomal proteins for phylogenomic analyses (Table 1). For bacterial dataset (B), the largest of the eight TQMD selections, this step took less than three hours to complete.

For each TQMD run, we assembled a supermatrix from the ribosomal proteins retrieved earlier (Table 1). Briefly, sequences were aligned with MAFFT v7.453 ^27^, then the alignments were cleaned using ali2phylip.pl from the Bio::MUST::Core software package (D. Baurain, https://metacpan.org/pod/Bio::MUST::Core), which implements the BMGE ^28^ filter (min=0.3, max=0.5, bmge-mask=loose). This step reduced the proportion of missing sites in the alignments. Next, we used Scafos v1.30 ^29^ to create the eight different supermatrices, using the Minimal evolutionary distance as a criterion for choosing sequences, the threshold set at 25%, the maximal percent of missing sites for a “complete sequence” set to 10 and the maximum number of missing OTUs set to 25, except for Firmicutes (22). Finally, IQ-TREE ^30,31^ was used to infer the phylogenomic tree associated with each supermatrix, using the LG4X model with ultrafast bootstraps. Trees were automatically annotated and colored using format-tree.pl (also from Bio::MUST::Core) and then visualised with iTOL v4 ^32^. The whole pipeline, from the launch of TQMD to the tree produced by IQ-TREE required approximately 3 days for the larger bacterial selection (Table 1, line B).

## Results and Discussion

The TQMD workflow has two separate phases: a preparation phase (Figure 1A) and a dereplication phase (Figure 1B). The objective of the preparation phase is to compute the genome-specific data that will be needed during the dereplication phase. These operations are embarrassingly parallel and very easy to speed up. In contrast, the dereplication phase considers all genomes at once, with the aim of clustering similar genomes based on pairwise distances and selecting the best representative for each cluster. To achieve this in the presence of many genomes, TQMD resorts to a greedy iterative heuristic in which each round is parallelized through a divide-and-conquer approach. The two phases are interconnected by the means of a relational database (see Materials and Methods for details). Hereafter, we study the effects of TQMD parameters and heuristics on its dereplication behavior, then we compare its performance to those of two similar solutions, dRep and Assembly Dereplicator and, finally, we provide some application examples in the field of bacterial phylogenomics.

### Analysis of TQMD behavior, parameters and heuristics

The dereplication phase is governed by a number of parameters and heuristics. One important issue is the inter-distance genome, which can either be the well-known Jaccard index (JI) or the identical genome fraction (IGF; see Materials and Methods for details). The latter was developed in an attempt to handle the comparison of genome pairs in which one is either partial or strongly reduced due to streamlining evolution ^20^. Whatever the selected distance, genomes that are less distant than a user-specified threshold will end up in the same cluster. This distance threshold is thus the main “knob” for controlling the aggressivity of TQMD dereplication: the lower the threshold the tighter the clustering. Another point to consider are TQMD heuristics and their parameterization. Since TQMD is iterative, one can always decide to dereplicate genomes that are themselves representatives obtained in one or more previous runs. When trying to dereplicate very large and taxonomically broad datasets, this raises the possibility to “guide” the dereplication by first clustering several phylum-wide subsets before merging the selected representatives in a single dataset to be dereplicated once more. This “indirect approach” is to be contrasted with the “direct approach”, in which TQMD is left dealing with the whole dataset from the very beginning. Regarding the divide-and-conquer algorithm operating during a single round, two parameters might be relevant: the pack size (e.g., 200 to 500) and the dividing scheme (random or taxonomically-guided). Obviously, larger pack sizes require more time to be processed but are less likely to be affected by the impossibility to dereplicate two genomes that are in different packs. Finally, in an attempt to balance such negative effects and the clustering speed, genome packs can either be composed at random (random sort) or by preferentially grouping taxonomically related organisms (taxonomic sort).

#### Performance criteria

Before studying the behavior of TQMD under different sets of parameters and heuristics, one has to recall that its aim is to generate dereplicated lists of genomes that maintain the phylogenetic diversity of the input genomes, especially at the highest levels of the prokaryotic taxonomy. Therefore, we identified two metrics of interest when examining TQMD output: (1) the number of phyla with at least one representative genome (“diversity”) and (2) the taxonomic mixing amongst the clusters (“mixity”). The diversity can be put in perspective with the number of representatives using what we call a redundancy index, i.e., the number of representatives divided by the number of phyla, with the lower the better. Regarding the concept of taxonomic mixing, we use it when the group of genomes behind a representative genome is not taxonomically homogeneous at some specific taxonomic level. Since our objective is mostly to dereplicate at the phylum level, we checked the taxonomic mixing at the phylum level. For example, if within a group of Proteobacteria, one (or several) Firmicutes is present, then the group is considered “mixed”.

#### Iterative algorithm: dereplication kinetics

We first compared the results of the two distance metrics (JI or IGF) on the full set of RefSeq Bacteria passing our quality control (see Materials and Methods). To study the effect of the distance threshold used for dereplication, we selected two ranges of six values giving similar final numbers of representatives for the two metrics (JI: from 0.1 to 0.2; IGF: from 0.3 to 0.4). Figure 3 shows the dereplication kinetics observed with the two metrics when using a medium threshold (JI: 0.16; IGF: 0.34) and the direct approach. The extreme efficacy (i.e., clustering ratio; see Materials and Methods) of the first round of dereplication is clear and subsequent rounds reach a plateau almost immediately. Whereas there is no notable difference between the two metrics in terms of kinetics, the height of the plateaus are not the same, with the IGF distance appearing greedier than the JI distance, especially when considering represented phyla rather than representative genomes.

**Figure 3.**
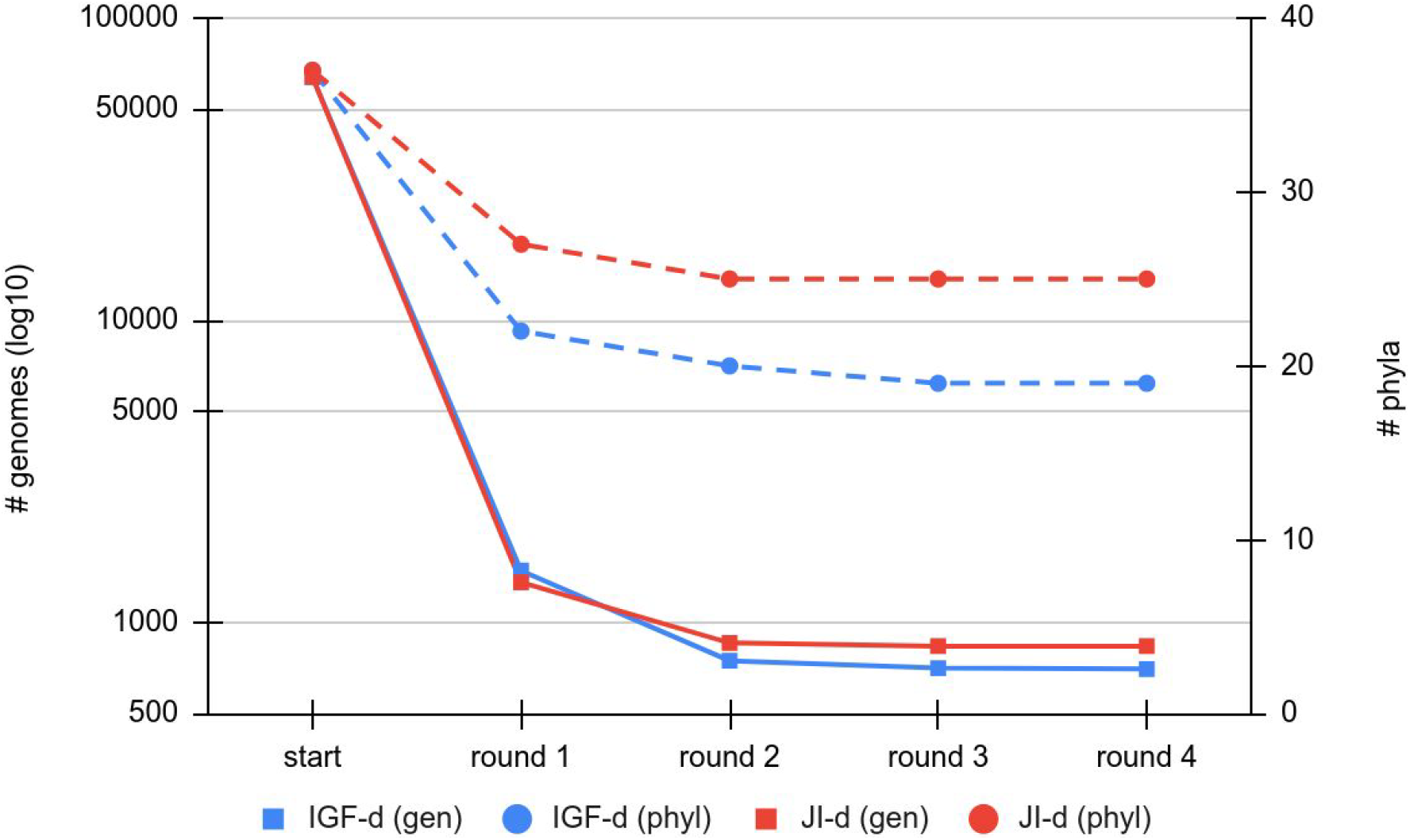
Comparison of the dereplication kinetics of TQMD when varying the distance metric. Two runs were launched on all RefSeq Bacteria (63,836 genomes; 37 phyla) using the direct approach and a pack size of 200, one with the Jaccard Index (JI-d, distance threshold of 0.16, red curves) and one with the Identical Genome Fraction (IGF-d, distance threshold of 0.34, blue curves). The left Y-axis shows the log10 of the number of remaining genomes (square dots and solids lines), whereas the right Y-axis shows the number of phyla for which at least one representative is still present at a given round of dereplication (round dots and dashed lines).

#### Iterative algorithm: effect of parameters and heuristics

While TQMD was designed to be run without manual intervention (direct approach), it is also possible to funnel the process by feeding it taxonomically homogeneous subsets of representative genomes (indirect approach). To contrast the two approaches, we first separated Bacteria into five groups corresponding to the four largest phyla in terms of numbers of genomes available in RefSeq and a fifth group with the rest of Bacteria: Proteobacteria (39,011 genomes), Firmicutes (26,972 genomes), Actinobacteria (10,248 genomes), Bacteroidetes (1639 genomes), other bacteria (2682 genomes).Then we dereplicated the four phyla separately using the Jaccard Index and a distance threshold of 0.2. Finally, we pooled the representatives obtained through the four TQMD runs with the remaining Bacteria and launched a final run on this reconstructed list. For this final run, we tried the two metrics and the full range of thresholds. The results of this multidimensional comparison are provided in Table 2 and Figure 4.

**Figure 4.**
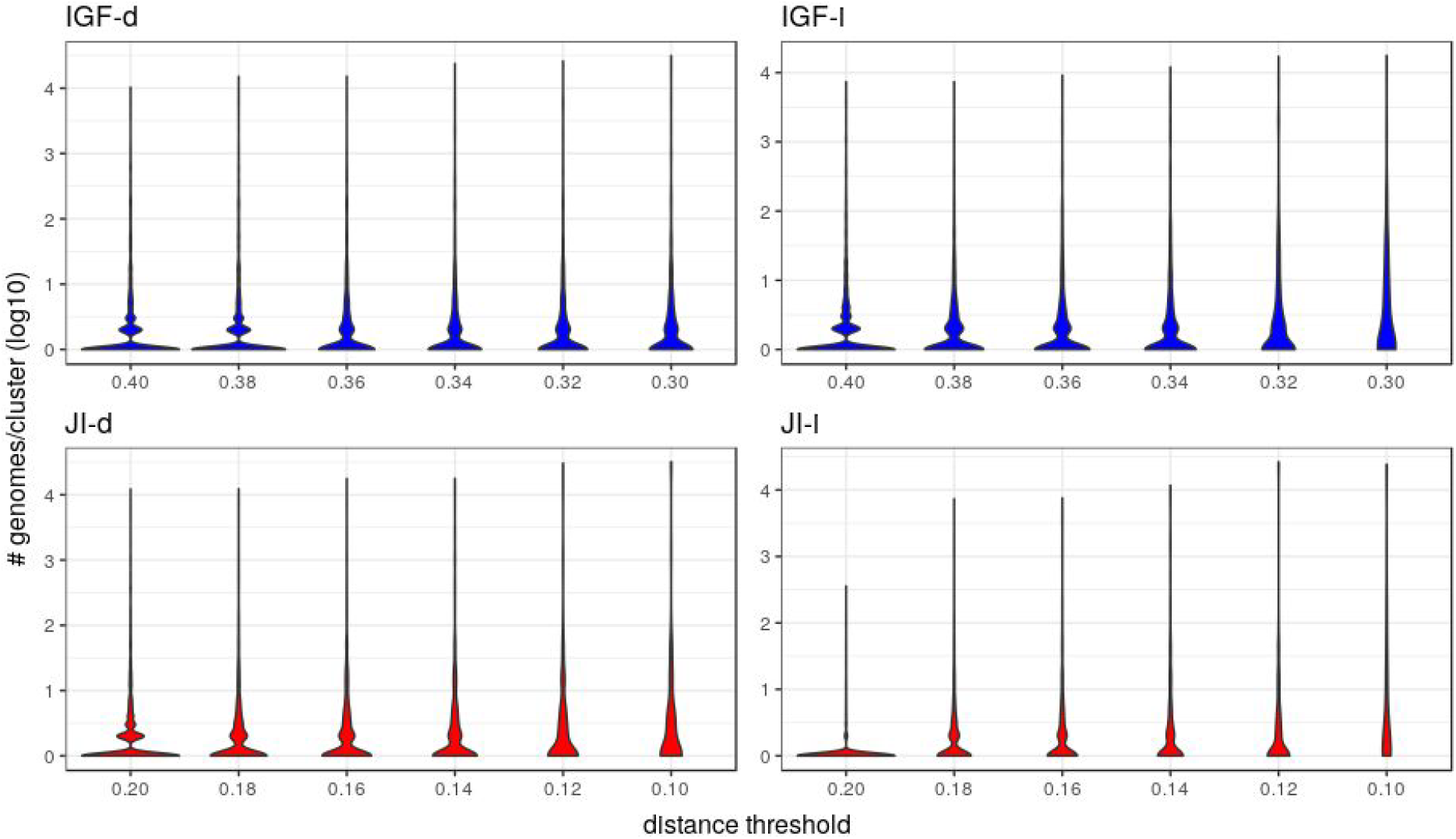
Distribution of the number of genomes per cluster when varying the distance metric, the distance threshold or the clustering approach. These violin plots are a companion to Table 2 and abbreviations are as in the latter table. The Y-axes are in log10 units and the violin plot width is proportional to the number of clusters containing the given number of genomes.

Starting with an initial number of bacterial phyla equal to 37, it appears that the two JI approaches are better than any IGF approach in terms of diversity, since the former retain a higher number of represented phyla for a given number of representative genomes. For example, when ending with about 500 representatives, the JI distance preserves 22-24 phyla, whereas the IGF distance only retains 15-19 phyla. These numbers translate to redundancy index (RI) values of 25-20 (JI) and 31-23 (IGF), respectively (Table 2). With the IGF distance, the indirect approach appears better at all thresholds, with a number of represented phyla systematically higher for a number of representatives systematically lower. This translates to, e.g., RI = 50 (IGF-i) vs 65 (IGF-d) with about 1550 representatives and RI = 30 (IGF-i) vs 33 (IGF-d) for about 720 representatives. In contrast, this is less obvious with the JI distance, where the indirect approach does not perform significantly better, the number of representatives also decreases but the number of represented phyla is also lower (or equal for the 0.1 threshold).

In the majority of the groups, the genome count per cluster is low with a significant proportion of singletons (i.e. only one representative genome, Table 2). However, in a few cases, large phyla (e.g., Proteobacteria, Firmicutes) gather into mixed groups that reach extreme genome counts and are visible as pikes in Figure 4. Neither approach changes this tendency but it is of notice that the JI distance with the indirect approach is the combination leading to the lowest genome count per cluster and the lowest count of mixed groups (Table 2 and panel JI-i in Figure 4), indicating a tendency to prevent the appearance of polyphyletic groups. When looking at the mixing (Table 2), it appears that unless at the lowest thresholds, the mixity remains marginal in all approaches. To analyze the situation within the mixed groups, we separated them into three categories: (1) paraphyletic groups (only one case, Firmicutes and Tenericutes), (2) super-groups (e.g., FBC, PVC, Terrabacteria; see Figure 6), and (3) polyphyletic groups. Since the TQMD objective is aggressive dereplication, the first two types of mixing are not problematic. Indeed they show that TQMD works as intended by first regrouping similar genomes together before regrouping the more dissimilar genomes. This also confirms that multiple scales of genuine phylogenetic signal lie in the nucleotide k-mers used in TQMD ^33,34^.

Amongst polyphyletic groups, the “early” groups, i.e., those that appear at higher thresholds (0.2 for JI and 0.4 for IGF), are (1) Firmicutes/Tenericutes clustered with Thermotogae and other thermophilic bacteria and (2) Terrabacteria clustered with Synergistetes. Thermotogae are likely mixed with Firmicutes due to their chimeric nature, Firmicutes being one of the main gene contributors (through lateral gene transfer, LGT) to Thermotogae ^35,36^. At lower thresholds, Thermotogae attract the other thermophilic bacteria, leading to the formation of a polyphyletic group. This result is a consequence of our single-linkage approach, which reveals to be a weakness when it comes to chimeric organisms that can bridge unrelated bacterial genomes. Regarding the clustering of Synergistetes with other Terrabacteria, when only a few genomes were available, Synergistetes were dispersed within two other phyla, Deferribacteres and Firmicutes ^37^. Nowadays, Synergistetes form a monophyletic group that is sister to Deferribacteres ^37^. We hypothesize that conflicting (maybe artifactual) signals cause (at least some) Synergistetes to cluster with Firmicutes, and then to attract other Terrabacteria in a snow-ball effect due to single linkage. In other words, as the thresholds are lowered, Thermotogae and Synergistetes serve as bridges between other bacterial phyla, creating or enlarging polyphyletic groups. This highlights that, just like alignment-based phylogeny, *k*-mer based approaches are also affected by chimeric organisms and LGT ^38^.

#### Divide-and-conquer algorithm: effect of parameters and heuristics

With respect to the parallelization of TQMD, the pack size has an influence on the results, since every time the size is diminished, the number of representatives returned at the end increases, whatever the distance metric (Table 3). This can be explained easily. In each pack, there is a list of genomes, to which each genome is compared in turn until it finds a cluster to join or creates a new cluster on its own. For each group, the selected representative is the best genome to work with in downstream applications, but not the “centroid” genome for the cluster. This means that a representative can be in the “outskirt” of its cluster in terms of sequence, which makes it less able to attract other genomes in subsequent groups. On the opposite, the single-linkage approach helps to alleviate the outskirt effect by enabling a genome to join a cluster as soon as any genome of that cluster is within the specified distance threshold. Another way to solve this issue is by increasing the pack size yet at the cost of speed. For example, 25 genomes require approximately 30 minutes to be processed, while 200 genomes take 2 hours and 500 genomes take several days, which corresponds to a quadratic complexity.

**Table 3.** Effect of the pack size on the final number of representative genomes. JI-based (direct) analyses were run using a distance threshold of 0.16, where IGF-based (direct) analyses used a threshold of 0.34. All of these analyses were run on 63,863 RefSeq Bacteria.

| pack size | # representatives |  |
| --- | --- | --- |
|  | JI-d | IGF-d |
| 200 | 836 | 702 |
| 100 | 874 | 866 |
| 50 | 966 | 1197 |
| 25 | 1041 | 1387 |

Finally, TQMD tries to speed up the dereplication process by assembling packs following a taxonomic sort of the genomes to dereplicate. This heuristic should improve the clustering ratio of each iteration by directly comparing genomes that are more likely to be similar, thereby greatly reducing the required number of rounds of the whole process. As expected, five independent runs launched on all RefSeq Bacteria using JI-d (Table 4) with genomes sorted randomly returned selections of 904 representatives (on average) in 17 to 18 rounds whereas, the same run with genomes sorted according to taxonomy returned 836 representatives in only four rounds. Similarly, five runs using IGF-d with genomes sorted randomly yielded 456 representatives (on average) in 9 to 10 rounds, in contrast to 702 representatives in four rounds by enabling the taxonomic sort. However, when dereplicating subsets corresponding to Proteobacteria, the random dividing scheme returned less representatives (124, worst result) than the taxonomic dividing scheme (165), in approximately the same number of rounds (3 to 5). Similar results were observed with Firmicutes: 224 representatives using the random scheme (worst result) vs 333 representatives using the taxonomic scheme. These results suggest that the random sort can be useful while working with a taxonomically homogeneous subset of bacteria. In other cases, it should be avoided because a higher number of rounds translates to a longer computing time.

**Table 4.** Comparison of the number of rounds and final representatives when modifying the distance metric and/or the dividing scheme for parallel processing. Five replicates of each combination were carried out for the random sort, whereas the taxonomic sort is deterministic. JI-based (direct) analyses were run using a distance threshold of 0.16, where IGF-based (direct) analyses used a threshold of 0.34. Pack size was 200.

| dataset | dist./appr. | sort | # rounds | # repr. |
| --- | --- | --- | --- | --- |
| Bacteria | JI-d | taxonomic | 4 | 836 |
| Bacteria | JI-d | random | 18 | 902 |
| Bacteria | JI-d | random | 17 | 903 |
| Bacteria | JI-d | random | 17 | 894 |
| Bacteria | JI-d | random | 18 | 915 |
| Bacteria | JI-d | random | 17 | 908 |
| Bacteria | IGF-d | taxonomic | 4 | 702 |
| Bacteria | IGF-d | random | 10 | 435 |
| Bacteria | IGF-d | random | 10 | 458 |
| Bacteria | IGF-d | random | 10 | 456 |
| Bacteria | IGF-d | random | 9 | 438 |
| Bacteria | IGF-d | random | 10 | 493 |
| Proteobacteria | IGF-d | taxonomic | 3 | 165 |
| Proteobacteria | IGF-d | random | 3 | 115 |
| Proteobacteria | IGF-d | random | 3 | 105 |
| Proteobacteria | IGF-d | random | 3 | 100 |
| Proteobacteria | IGF-d | random | 3 | 124 |
| Proteobacteria | IGF-d | random | 3 | 114 |
| Firmicutes | IGF-d | taxonomic | 4 | 333 |
| Firmicutes | IGF-d | random | 4 | 190 |
| Firmicutes | IGF-d | random | 5 | 212 |
| Firmicutes | IGF-d | random | 5 | 224 |
| Firmicutes | IGF-d | random | 4 | 194 |
| Firmicutes | IGF-d | random | 4 | 172 |

### Comparison with dRep and Assembly-Dereplicator

When we began our work on TQMD in 2015, there was no published program for genome dereplication. Now two different software packages are available, dRep ^39^ and Assembly-Dereplicator [R. R. Wick and K. E. Holt, unpublished; https://github.com/rrwick/Assembly-Dereplicator], both built on top of Mash ^40^. Mash itself was created to estimate the Jaccard distance within sets of genomes and metagenomes based on nucleotide *k*-mer counts ^40^. dRep was designed especially for the dereplication of metagenomes, whereas Assembly-Dereplicator (A-D) was designed for groups of bacteria which are sufficiently close relatives. A comparison of the working principles of dRep, A-D and TQMD is available in Table 5.

**Table 5.** Feature comparison between dRep, Assembly-Dereplicator (A-D) and TQMD. JI: Jaccard Index, IGF: Identical Genome Fraction, ANI: average nucleotide identity, d-and-c: divide-and-conquer, SGE/OGE: Sun/Open Grid Engine, Y: present feature, N: absent feature.

| feature | dRep | A-D | TQMD |
| --- | --- | --- | --- |
| main engine(s) | Mash + ANIm (or gANI) | Mash | JELLYFISH |
| other dependencies | CheckM (optional) | none | QUAST, RNAmmer, Forty-Two (optional) |
| relational database | N | N | Y |
| distance metric(s) | JI (estimated) then ANI | JI (estimated) | JI (exact) or IGF (exact) |
| heuristic(s) | biphasic approach: Mash for fast and rough clustering followed by ANI for slow and accurate clustering | d-and-c strategy (serial) | iterative greedy algorithm (serial) + d-and-c strategy (parallel) |
| stop condition(s) | unspecified | first failure to dereplicate any serial batch | any of 3 possible cut-offs (number of rounds, number of representatives, clustering ratio) |
| d-and-c dividing scheme | unspecified | random | random or taxonomic |
| selection of representatives | formula based on genome size, assembly quality and contamination level (incl. strain heterogeneity) | assembly quality | formula based on genome size, assembly quality, annotation richness and contamination level |
| parameterization of representative selection | Y (parameter weights) | N | Y (simplified formula) |
| automatic genome download | N | N | Y |
| grid engine support | N | N | Y (SGE/OGE) |
| CPU use | fixed on launch | fixed on launch | specified as a maximum (decreases over time) |

To compare TQMD to dRep (v2.2.3), we chose two different datasets, the phylum Bacteroidetes (1127 genomes) and the order Streptomycetales (648 genomes; phylum Actinobacteria). Because of technical difficulties with the installation of dRep, we had to use a workstation less powerful than the grid computer used to run TQMD (see Materials and Methods). That is why we did not use all the available bacterial genomes in these tests. Regarding Bacteroidetes, dRep required five hours (using 10 CPUs and default parameters) to select 835 genomes. With TQMD, we used a threshold of 0.4 on the Jaccard distance to obtain comparable results. TQMD run lasted 10 hours (on at most 6 CPUs) and selected 789 representative genomes, of which 707 were in common with those of dRep. Since our main objective is to maintain as much as possible the diversity when dereplicating, we verified how many species were retained after the dereplication. Before dereplication, we had 528 different species of Bacteroidetes; dRep produced a list covering 516 of these species, whereas TQMD produced a list of 517 species, of which 511 were in common (see table 6 for details). With Streptomycetales, dRep (again using default values), selected 430 genomes out of 648 in approximately 12h30min using 20 CPUs. To emulate such a result with TQMD, we had to use a threshold of 0.6 and obtained 486 representatives (392 in common, of which 39 species, 342 corresponding to organisms not identified down to the species) in about 10h using at most 4 CPUs in parallel (details given in Table 6).

**Table 6.** Performance comparison between TQMD and dRep on two smaller datasets. # gen.: starting number of genomes, # repr.: final/common number of representative genomes, # spec. starting/final/common number of species, h.CPU: upper bound on CPU use (i.e., product of wall-clock time and number of CPUs). (*) Out of 392 genomes, 342 are not identified down to the species (e.g., *Streptomyces* sp. GCF_000470535.1). With TQMD, a distance threshold of 0.4 was used for Bacteroidetes and a threshold of 0.6 for Streptomycetales. The pack size was 200 and a taxonomic sort was used for both cases.

| dataset | starting |  | TQMD |  |  | dRep |  |  | intersection |  |
| --- | --- | --- | --- | --- | --- | --- | --- | --- | --- | --- |
|  | # gen. | # spec. | # repr. | # spec. | h.CPU | # repr. | # spec. | h.CPU | # repr. | # spec. |
| Bacteroidetes | 1127 | 528 | 789 | 517 | 60 | 835 | 516 | 50 | 707 | 511 |
| Streptomycetales | 648 | 220 | 486 | 207 | 40 | 430 | 189 | 250 | 392 | 39* |

dRep is a less aggressive program than TQMD, which is unsurprising as the former is meant to be used on sets of metagenomes and to dereplicate at the species level, while the latter is meant to be used on every completely sequenced prokaryotic genome available and to dereplicate at the phyla/class level. Moreover, from the very start, TQMD was designed with scalability in mind, so as to accommodate the ever growing number of sequenced genomes. In principle, dRep could be used aggressively like TQMD, by fine-tuning two different thresholds (primary and secondary clusters), but this would need dRep to allow the user to choose a different Mash *k*-mer size, which does not appear to be possible (for the average user). On the other hand, TQMD can be used to dereplicate down to the species level more easily (only one threshold to specify) but it would take a longer time to finish since it would require a longer JELLYFISH *k*-mer size (see Material and Methods). In conclusion, the dRep and TQMD can do each other’s work but become less efficient when trying to do so, thereby rather making them complementary: dRep to dereplicate at the species level and TQMD at phylum/class level. For intermediate taxonomic levels, it is up to the user to decide which one s/he prefers.

Assembly-Dereplicator is a program that is more recent but, as of August 2020, not yet published; its last update dates from November 2019. Its main advantage is ease of use, since it is a simple (no-installation) script that only needs Mash as a prerequisite. A-D takes as input the path to a folder containing the genomes to be dereplicated and rearranges them randomly and separated into smaller packs (500 genomes per pack by default). The next step is the clustering of each pack serially using Mash. A-D stops as soon as it cannot dereplicate at least one genome from the current pack. Since it was compatible with our grid computer, we tried to test A-D (v0.1.0) with all prokaryotic RefSeq genomes, so as to mimic how TQMD is supposed to work. In March 2019, this amounted to 112,254 genomes. With a threshold of 0.01 (i.e., two genomes are clustered if 99% of their nucleotides are identical) A-D stopped after one hour, only dereplicating three groups of 500 genomes to yield 111,855 “representative” genomes in output. With a more lenient threshold (0.1), the program behaved quite similarly, this time dereplicating nine groups of 500 genomes (110,160 genomes in output). We kept trying, raising the threshold by increments of 0.1 until we reached 0.5. Every time, the program stopped before completion and the results revealed to be disappointing. For example, at 0.5 (clustered if 50% of the genomes are identical), it worked on the first cluster then stopped at the second, yielding 111,755 genomes in output (details given in Table S1).

Apparently, the A-D script does not work with very large and non-homogeneous groups of genomes, probably due to the heuristics used. We did not go further investigating the script since our tests clearly showed that it cannot be compared, in its present state, with TQMD on very large datasets. Albeit A-D seems promising to dereplicate smaller sets of homogeneous genomes (such as the Cyanobacteria), exploring the efficiency of A-D on small datasets was not the purpose of this article. Yet, drawing on our own tests with TQMD (see above), our intuition is that the A-D approach based on a random splitting of the genomes to dereplicate, if appropriate when working with homogeneous genomes, fails when it comes to non-homogeneous datasets. Moreover, the stop conditions of the iterative heuristics obviously lead A-D to get stuck very easily.

### Application example of TQMD

To check whether TQMD output was indeed useful in a practical context, we computed phylogenomic trees based on concatenations of ribosomal proteins sampled from selected representative genomes. We performed two runs on all RefSeq Bacteria (63,863 genomes passing our prerequisites ; see Materials and Methods for details) using the indirect approach and the Jaccard Index, one at a distance threshold of 0.1 (Table 1, line A) and the other at 0.12 (Table 1, line B). The first run yielded a selection of 49 genomes while the second run retained 151 genomes. Six additional runs using the direct approach were carried out on the six largest bacterial phyla of RefSeq (in terms of numbers of organisms): Proteobacteria, Firmicutes, Actinobacteria, Bacteroidetes, Cyanobacteria and Chlamydia. These phylum-wide selections contained about 20 to 50 genomes, each collectively representing the diversity of their respective phyla (Table 1, line C-H). In this text, we only show and describe the larger phylogenomic tree of all Bacteria (Table 1, line B). The seven other trees are available as Figures S1 to S7.

The larger bacterial tree (Figure 5) results from an extremely aggressive selection (Table 1, B) but it still shows what we consider as the main groups of Bacteria (Proteobacteria, PVC, FBC, and monoderms phyla) and, after accounting for the idiosyncratic taxon names, most groups described by T. Cavalier-Smith ^41^ are visible (with the exception of Eoglycobacteria and Hadobacteria, which were both absorbed in polyphyletic groups).

**Figure 5:**
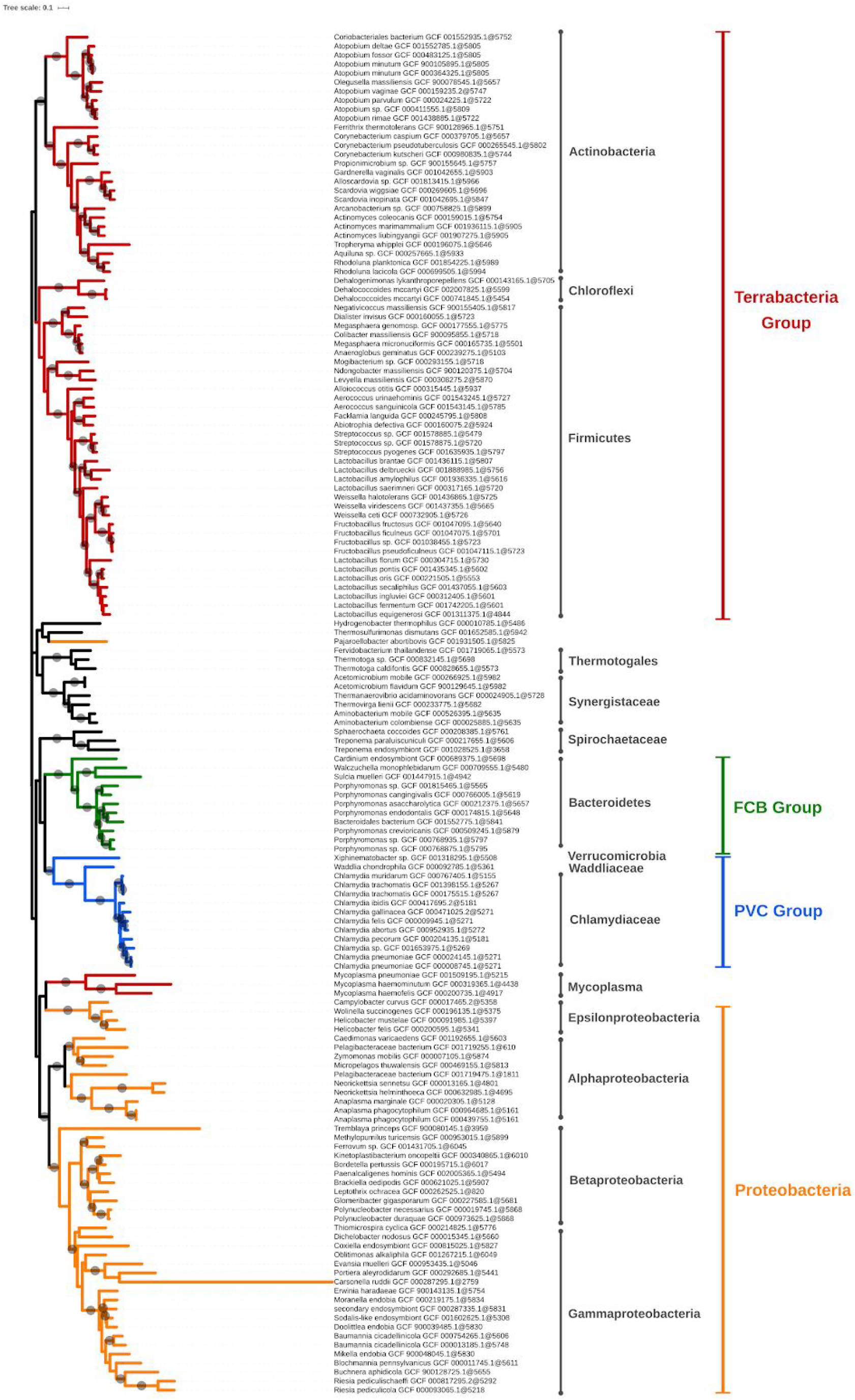
Phylogenetic tree of the largest selection of Bacteria. (Table 1, B) inferred from a supermatrix of concatenated ribosomal proteins under the LG4X model using IQ-TREE. Dots on branches indicate maximum bootstrap support values (100%).

Regarding the topology of the tree, all the organisms from the main super-phyla are generally regrouped in the same subtree, with some exceptions. These exceptions are the mycoplasma branch, which ends up within Proteobacteria, and *Pajaroellobacter abortibovis*, a proteobacterium that is separated from other Proteobacteria.

In Figure 5, some genera and even species appear to be overrepresented in the selected genomes and form monophyletic subtrees within the tree. This is the case of *Lactobacillus*, for example, with 11 representatives (10 species). To investigate an eventual selection bias in TQMD, we launched two different TQMD runs using only the *Lactobacillus* genomes (841 which passed TQMD prerequisites). Both runs used the same values as the larger run for Bacteria (Table 1, B). The difference was the way of sorting the genomes before dividing them in packs, one used the taxonomic sort and the other the random sort. The run with the taxonomic sort yielded 19 *Lactobacillus* representatives (15 species), of which 10 in common with the larger run for Bacteria, whereas the random sort run yielded 21 representatives (16 species), of which 10 in common with the larger run for Bacteria and 16 with the taxonomic run. These results suggest that the taxonomic sort does not especially lead to a selection biased towards identically named genera or species, but that the representative genomes adequately sample the underlying phylogenetic diversity of the group. Along the same line, dRep results for Bacteroidetes also show genomes of the same “species” not clustered together as in our Bacteroidetes tree (Table 6 and Figure S3). This indicates that the genomes of such identically named organisms are actually quite different, thereby not reflecting a technical issue of TQMD or of dRep, but rather a genuine property of these genomes. Consequently, it is worth mentioning that a purely taxonomic (i.e., manual) selection of representative genomes would have overlooked this genomic diversity, thereby reducing the relevance of the selection.

## Conclusion

TQMD is an efficient dereplication tool designed for the assembly of phylum-level datasets of representative prokaryotic genomes. It manages to maintain the taxonomic diversity of input genomes while being fast, owing to its aggressive dereplication heuristics, which makes it able to scale with the ever growing number of genome assemblies in public repositories, such as NCBI RefSeq. At lower taxonomic levels, TQMD becomes slower, probably because it has to compare more genomes before finding pairs close enough to be clustered and dereplicated. It would also require to use a longer *k*-mer size in order to improve the dereplication, slowering TQMD further. In these cases, it is advised to use dRep, which is already optimized for these taxonomic levels. The development of the first version of TQMD is finished but could be improved notably by adding new distance metrics beyond JI and IGF, and/or by including additional variables for the selection of representative genomes. The two other programs considered here both rely on Mash instead of Jellyfish in our case. A revision of TQMD replacing Jellyfish by Mash would be interesting to compare the efficiency with the current version but this would imply dropping the IGF distance (since Mash only implements JI) and would require Mash to accept using different *k*-mers sizes.

## Authors’ contribution

DB and FK conceived the study. RRL developed ToRQuEMaDA and performed all analyses except phylogenomic analyses, which were carried out by ML with the help of MVV. RRL and DB wrote the manuscript. All authors read and approved the final manuscript.

## Acknowledgment

RRL and MVV were supported by FRIA fellowships of the Belgian National Fund for Scientific Research (F.R.S.-FNRS). FK is a Research Associate of the F.R.S.-FNRS. ML is supported by the French Agence Nationale de la Recherche (ANR, project MATHTEST). Computational resources were provided through two grants to Denis Baurain (University of Liège “Crédit de démarrage 2012” SFRD-12/04; F.R.S.-FNRS “Crédit de recherche 2014” CDR J.0080.15). The authors are grateful to Luc Cornet for his help in various areas of the work.

---

**Supplementary Table 1.**
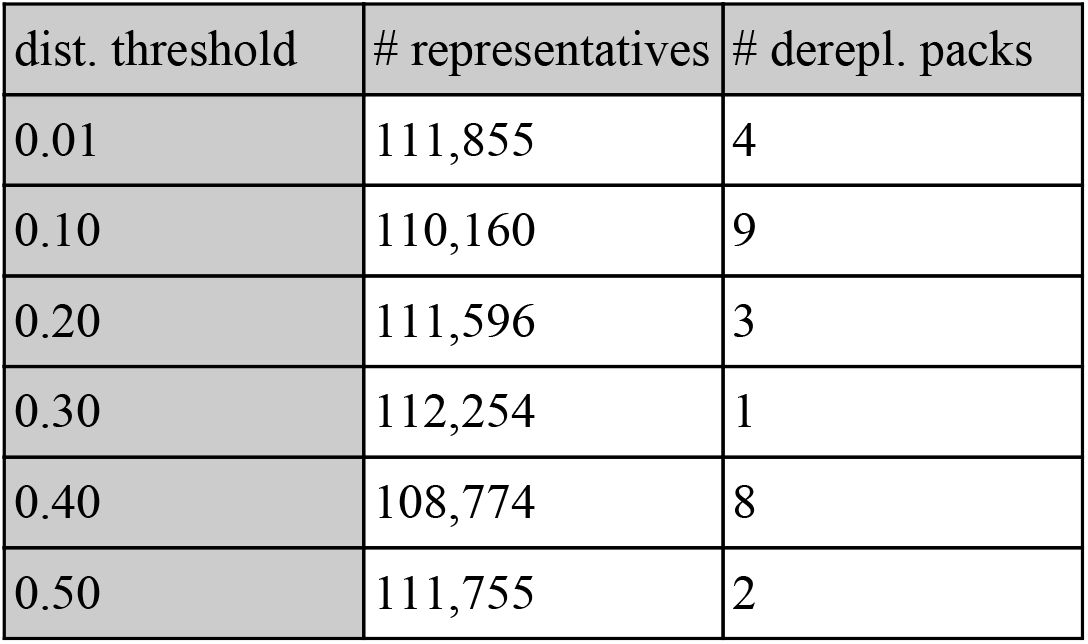
Attempts at dereplicating all RefSeq Bacteria (releases 79+92, 112,254 genomes) using Assembly-Dereplicator. Analyses were run using 6 different distance thresholds and the default pack size of 500 (225 packs).

| dist. threshold | # representatives | # derepl. packs |
| --- | --- | --- |
| 0.01 | 111,855 | 4 |
| 0.10 | 110,160 | 9 |
| 0.20 | 111,596 | 3 |
| 0.30 | 112,254 | 1 |
| 0.40 | 108,774 | 8 |
| 0.50 | 111,755 | 2 |

**Supplementary Figure 1:**
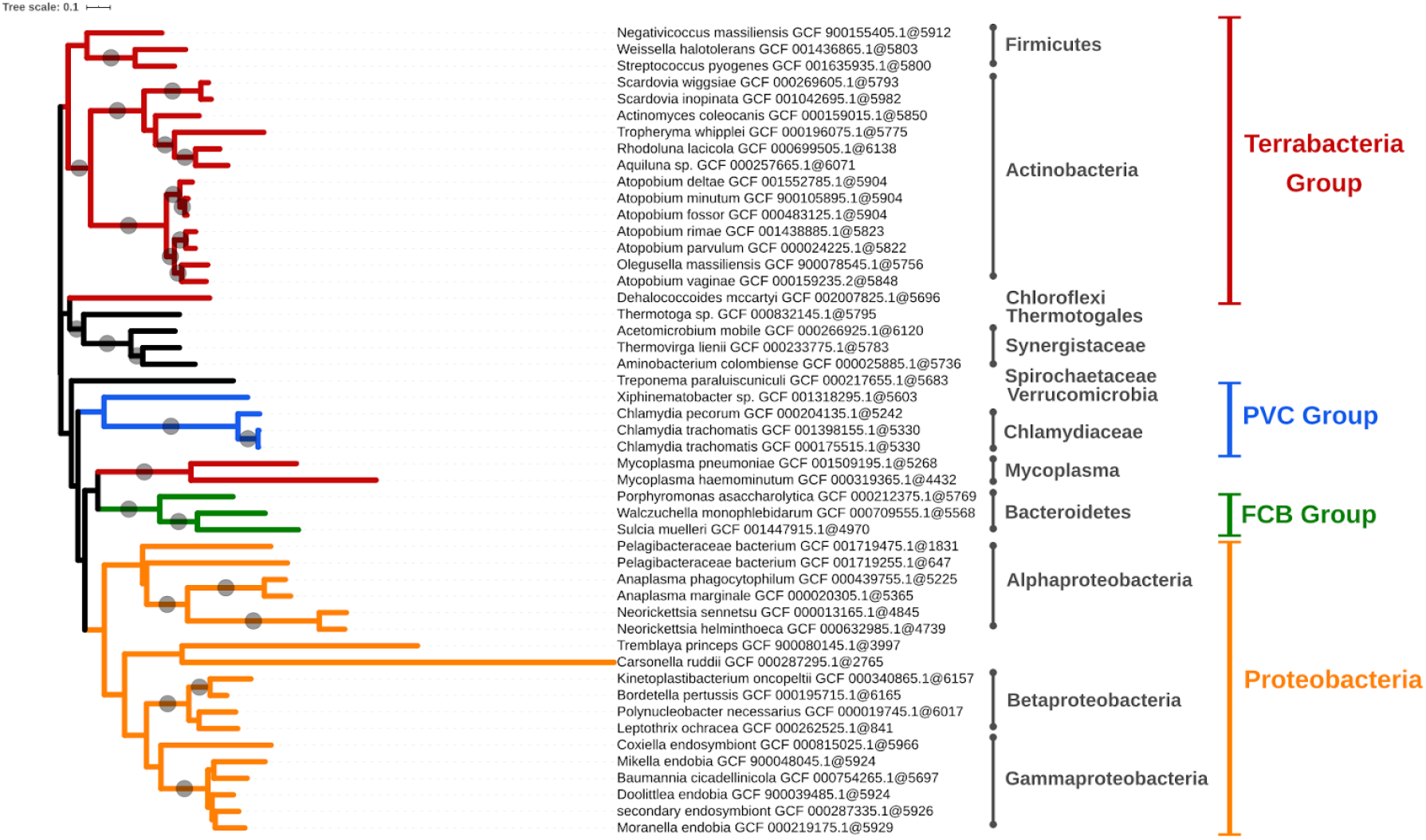
Phylogenetic tree of the smallest selection of Bacteria. (Table 1, A) inferred from a supermatrix of concatenated ribosomal proteins under the LG4X model using IQ-TREE. Dots on branches indicate maximum bootstrap support values (100%).

**Supplementary Figure 2:**
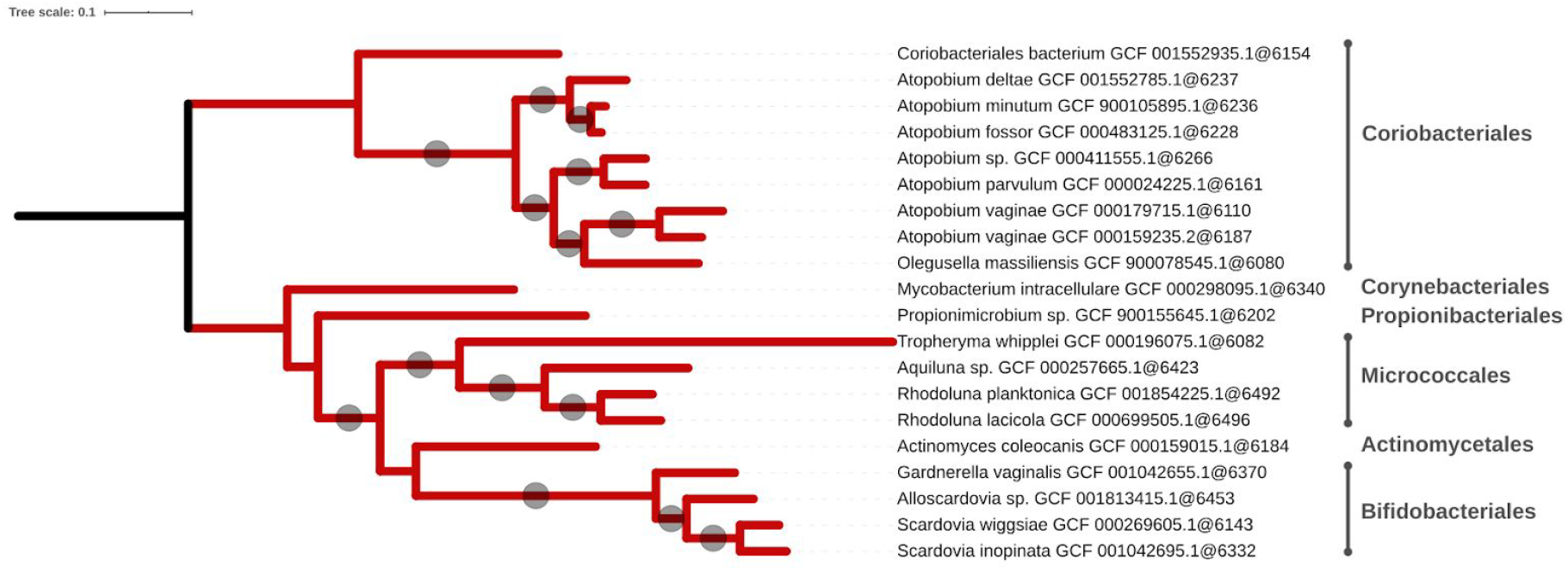
Phylogenetic tree of the Actinobacteria. (Table 1, C) inferred from a supermatrix of concatenated ribosomal proteins under the LG4X model using IQ-TREE. Dots on branches indicate maximum bootstrap support values (100%).

**Supplementary Figure 3:**
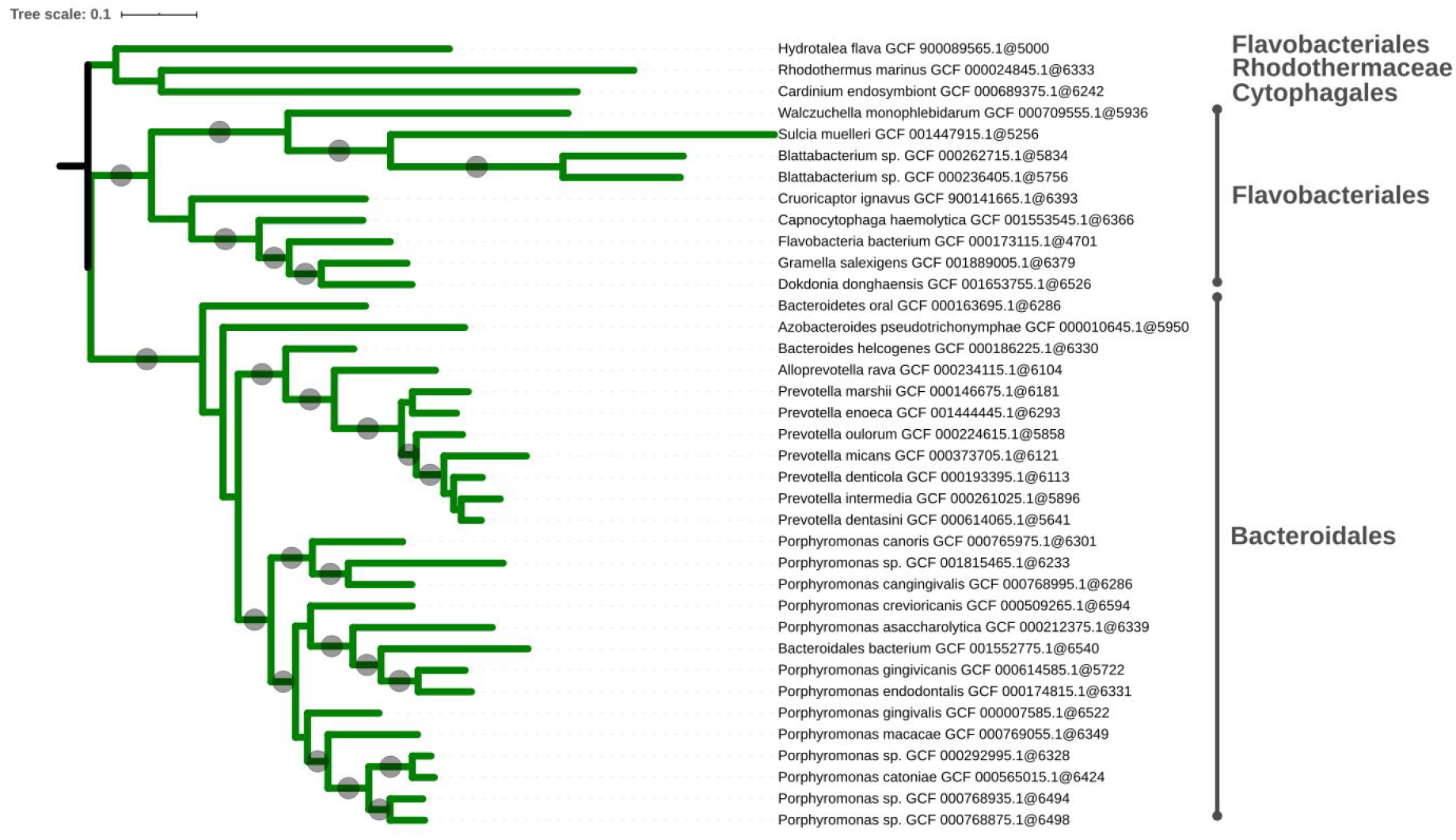
Phylogenetic tree of the Bacteroidetes. (Table 1, D) inferred from a supermatrix of concatenated ribosomal proteins under the LG4X model using IQ-TREE. Dots on branches indicate maximum bootstrap support values (100%).

**Supplementary Figure 4:**
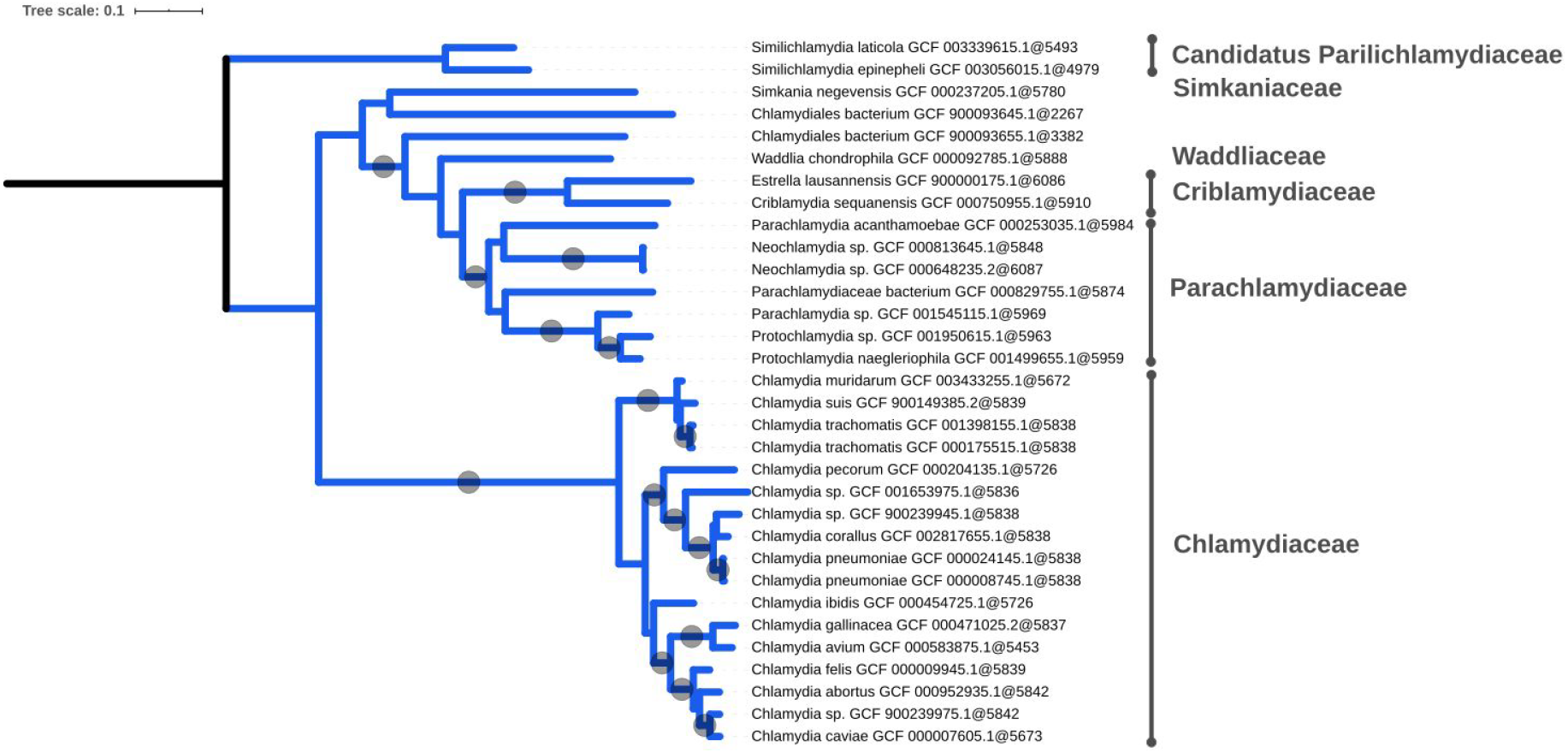
Phylogenetic tree of the Chlamydia. (Table 1, E) inferred from a supermatrix of concatenated ribosomal proteins under the LG4X model using IQ-TREE. Dots on branches indicate maximum bootstrap support values (100%).

**Supplementary Figure 5:**
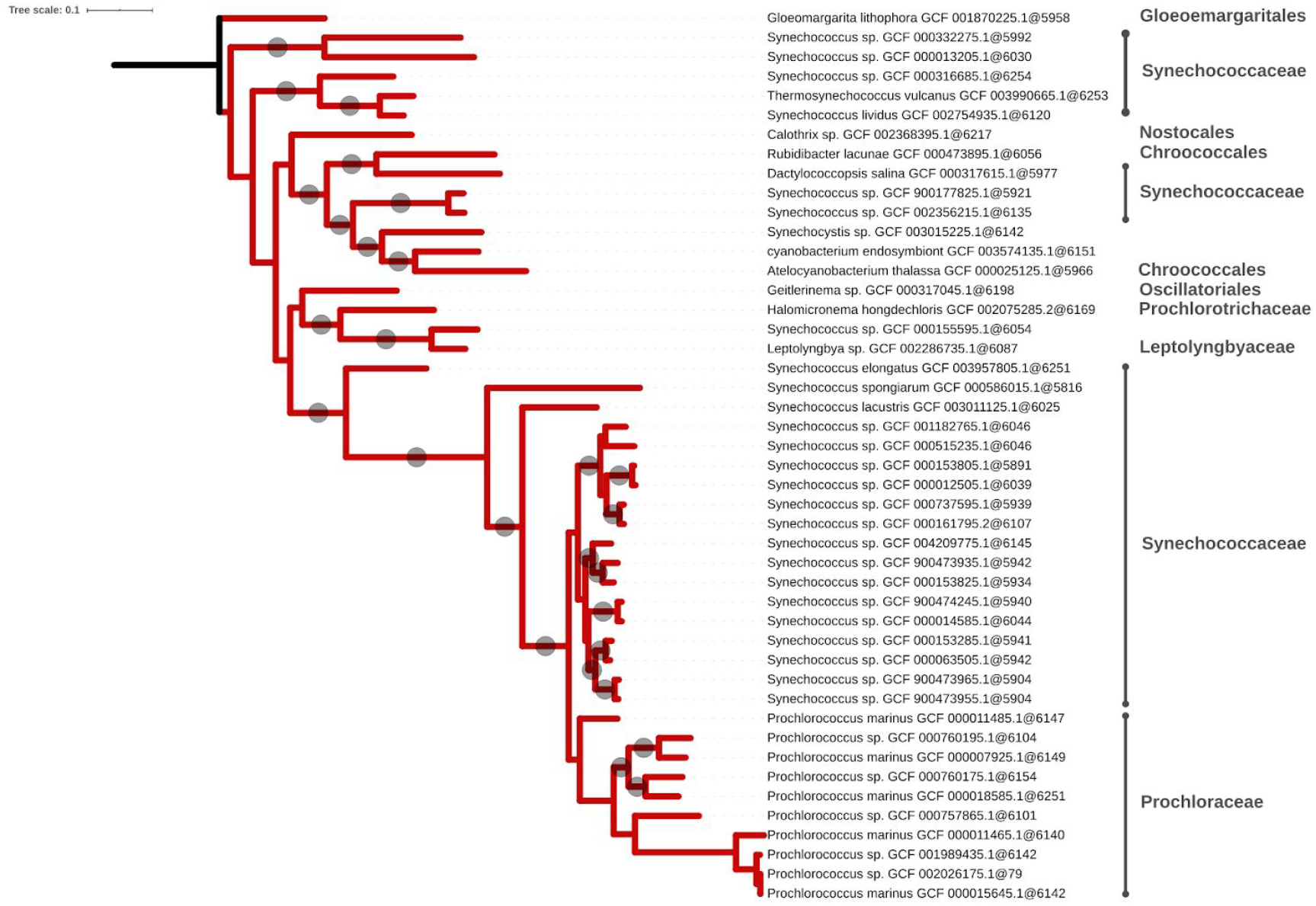
Phylogenetic tree of the Cyanobacteria. (Table 1, F) inferred from a supermatrix of concatenated ribosomal proteins under the LG4X model using IQ-TREE. Dots on branches indicate maximum bootstrap support values (100%).

**Supplementary Figure 6:**
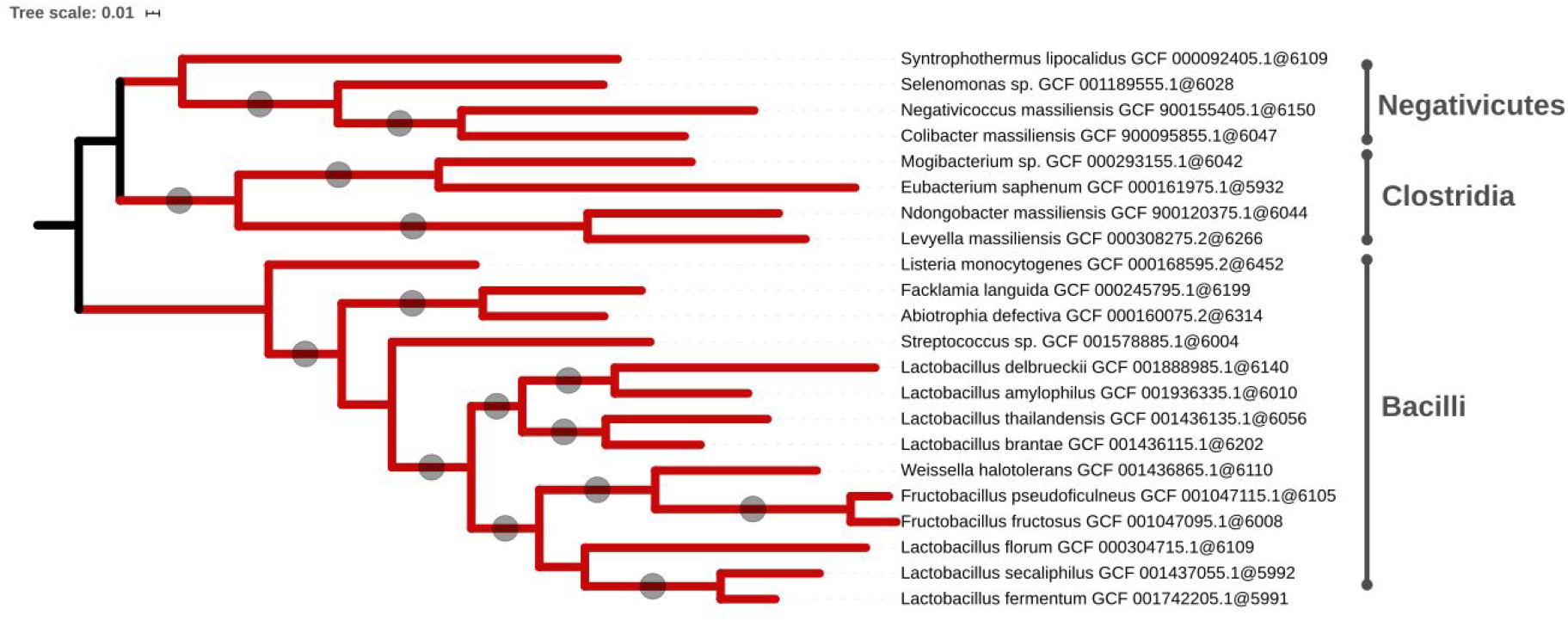
Phylogenetic tree of the Firmicutes. (Table 1, G) inferred from a supermatrix of concatenated ribosomal proteins under the LG4X model using IQ-TREE. Dots on branches indicate maximum bootstrap support values (100%).

**Supplementary Figure 7:**
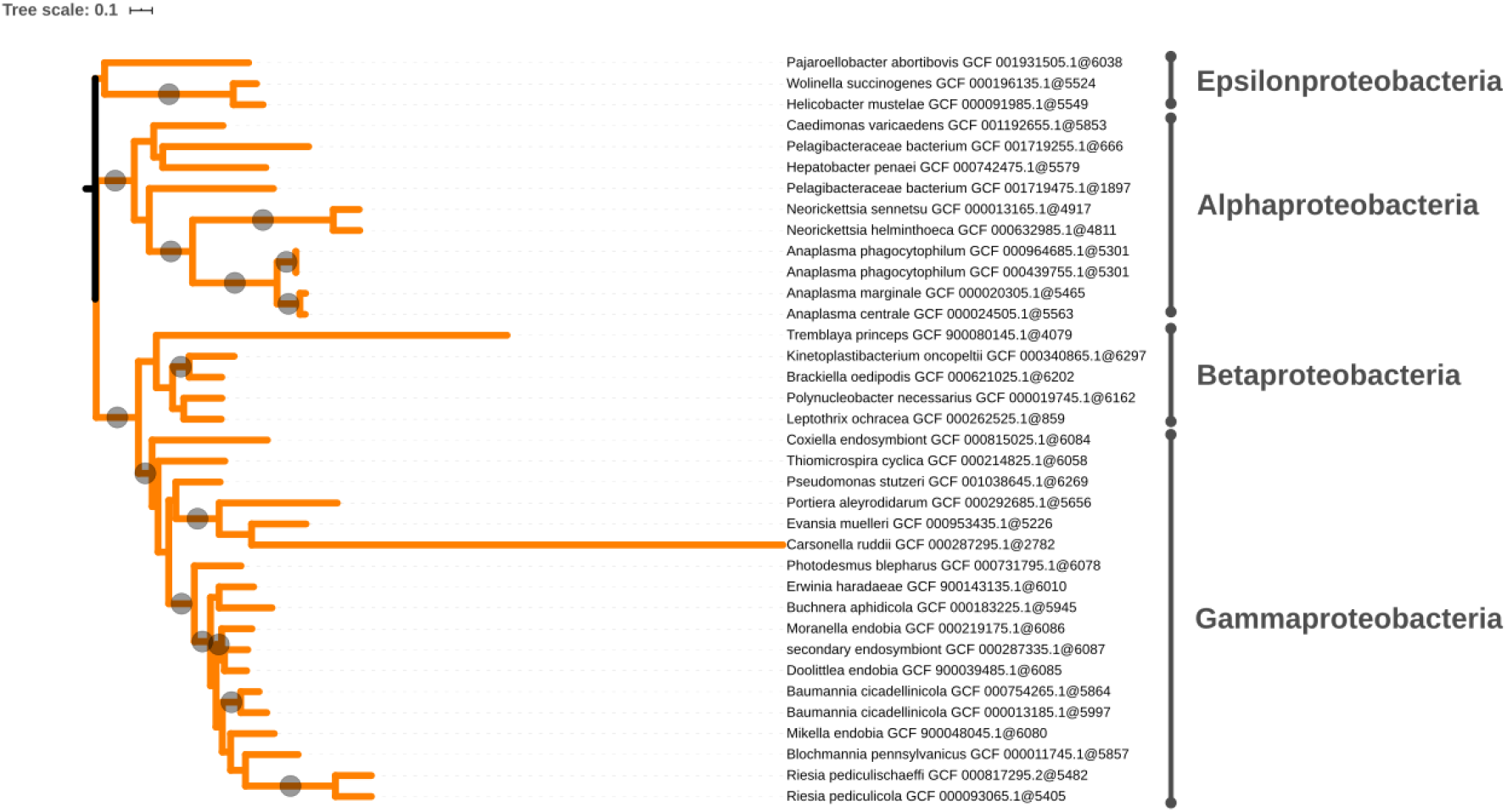
Phylogenetic tree of the Proteobacteria. (Table 1, H) inferred from a supermatrix of concatenated ribosomal proteins under the LG4X model using IQ-TREE. Dots on branches indicate maximum bootstrap support values (100%).

## References

1. Zielezinski, A., Vinga, S., Almeida, J. & Karlowski, W. M. Alignment-free sequence comparison: benefits, applications, and tools. Genome Biol. 18, 186 (2017).

2. Shannon, C. E. lJ A mathematical theory of communication. Bell System Tech. J. 27, 379-423, 623-656 (1948).-[2. Certain Results Coding Theory Noisy Channels Inf. Controll 6–25 (1957).

3. Kullback, S. & Leibler, R. A. On information and sufficiency. Ann. Math. Stat. 22, 79–86 (1951).

4. Tribus, M. & McIrvine, E. C. Energy and information. Sci. Am. 225, 179–190 (1971).

5. Batista, M. V. A., Ferreira, T. A. E., Freitas, A. C. & Balbino, V. Q. An entropy-based approach for the identification of phylogenetically informative genomic regions of Papillomavirus. Infect. Genet. Evol. 11, 2026–2033 (2011).

6. Kolmogorov, A. N. Three approaches to the quantitative definition ofinformation’. Probl. Inf. Transm. 1, 1–7 (1965).

7. Chan, C. X., Bernard, G., Poirion, O., Hogan, J. M. & Ragan, M. A. Inferring phylogenies of evolving sequences without multiple sequence alignment. Sci. Rep. 4, 6504 (2014).

8. Bowers, R. M. et al. Minimum information about a single amplified genome (MISAG) and a metagenome-assembled genome (MIMAG) of bacteria and archaea. Nat. Biotechnol. 35, 725–731 (2017).

9. Irisarri, I. et al. Phylotranscriptomic consolidation of the jawed vertebrate timetree. Nat. Ecol. Evol. 1, 1370–1378 (2017).

10. Simion, P. et al. A large and consistent phylogenomic dataset supports sponges as the sister group to all other animals. Curr. Biol. 27, 958–967 (2017).

11. O’Leary, N. A. et al. Reference sequence (RefSeq) database at NCBI: current status, taxonomic expansion, and functional annotation. Nucleic Acids Res. 44, D733–D745 (2016).

12. Marçais, G. & Kingsford, C. A fast, lock-free approach for efficient parallel counting of occurrences of k-mers. Bioinformatics 27, 764–770 (2011).

13. Gurevich, A., Saveliev, V., Vyahhi, N. & Tesler, G. QUAST: quality assessment tool for genome assemblies. Bioinformatics 29, 1072–1075 (2013).

14. Cornet, L. et al. Consensus assessment of the contamination level of publicly available cyanobacterial genomes. PloS One 13, e0200323 (2018).

15. Lagesen, K. et al. RNAmmer: consistent and rapid annotation of ribosomal RNA genes. Nucleic Acids Res. 35, 3100–3108 (2007).

16. Li, W. & Godzik, A. Cd-hit: a fast program for clustering and comparing large sets of protein or nucleotide sequences. Bioinformatics 22, 1658–1659 (2006).

17. Fu, L., Niu, B., Zhu, Z., Wu, S. & Li, W. CD-HIT: accelerated for clustering the next-generation sequencing data. Bioinformatics 28, 3150–3152 (2012).

18. Taton, A., Grubisic, S., Brambilla, E., De Wit, R. & Wilmotte, A. Cyanobacterial diversity in natural and artificial microbial mats of Lake Fryxell (McMurdo Dry Valleys, Antarctica): a morphological and molecular approach. Appl. Environ. Microbiol. 69, 5157–5169 (2003).

19. Edgar, R. C. Updating the 97% identity threshold for 16S ribosomal RNA OTUs. Bioinformatics 34, 2371–2375 (2018).

20. Cornet, L. et al. Metagenomic assembly of new (sub) polar Cyanobacteria and their associated microbiome from non-axenic cultures. Microb. Genomics 4, (2018).

21. Jauffrit, F. et al. RiboDB database: a comprehensive resource for prokaryotic systematics. Mol. Biol. Evol. 33, 2170–2172 (2016).

22. Real, R. & Vargas, J. M. The Probabilistic Basis of Jaccard’s Index of Similarity. 45, 380–385 (1996).

23. Jones, N. C., Pevzner, P. A. & Pevzner, P. An introduction to bioinformatics algorithms. (MIT press, 2004).

24. Bentley, J. L. Multidimensional divide-and-conquer. Commun. ACM 23, 214–229 (1980).

25. Federhen, S. The NCBI taxonomy database. Nucleic Acids Res. 40, D136–D143 (2012).

26. Sayers, E. W. et al. GenBank. Nucleic Acids Res. 48, D84–D86 (2020).

27. Katoh, K. & Standley, D. M. MAFFT multiple sequence alignment software version 7: improvements in performance and usability. Mol. Biol. Evol. 30, 772–780 (2013).

28. Criscuolo, A. & Gribaldo, S. BMGE (Block Mapping and Gathering with Entropy): a new software for selection of phylogenetic informative regions from multiple sequence alignments. BMC Evol. Biol. 10, 210 (2010).

29. Roure, B., Rodriguez-Ezpeleta, N. & Philippe, H. SCaFoS: a tool for selection, concatenation and fusion of sequences for phylogenomics. BMC Evol. Biol. 7 Suppl 1, S2 (2007).

30. Nguyen, L.-T., Schmidt, H. A., Von Haeseler, A. & Minh, B. Q. IQ-TREE: a fast and effective stochastic algorithm for estimating maximum-likelihood phylogenies. Mol. Biol. Evol. 32, 268–274 (2015).

31. Hoang, D. T., Chernomor, O., Von Haeseler, A., Minh, B. Q. & Vinh, L. S. UFBoot2: improving the ultrafast bootstrap approximation. Mol. Biol. Evol. 35, 518–522 (2018).

32. Letunic, I. & Bork, P. Interactive Tree Of Life (iTOL) v4: recent updates and new developments. Nucleic Acids Res. 47, W256–W259 (2019).

33. Wen, J., Chan, R. H., Yau, S.-C., He, R. L. & Yau, S. S. K-mer natural vector and its application to the phylogenetic analysis of genetic sequences. Gene 546, 25–34 (2014).

34. Allman, E. S., Rhodes, J. A. & Sullivant, S. Statistically consistent k-mer methods for phylogenetic tree reconstruction. J. Comput. Biol. 24, 153–171 (2017).

35. Nesbø, C. L. et al. The genome of Thermosipho africanus TCF52B: lateral genetic connections to the Firmicutes and Archaea. J. Bacteriol. 191, 1974–1978 (2009).

36. Gupta, R. S. & Bhandari, V. Phylogeny and molecular signatures for the phylum Thermotogae and its subgroups. Antonie Van Leeuwenhoek 100, 1 (2011).

37. Jumas-Bilak, E., Roudiere, L. & Marchandin, H. Description of ‘Synergistetes’ phyl. nov. and emended description of the phylum ‘Deferribacteres’ and of the family Syntrophomonadaceae, phylum ‘Firmicutes’. Int. J. Syst. Evol. Microbiol. 59, 1028–1035 (2009).

38. Daubin, V., Moran, N. A. & Ochman, H. Phylogenetics and the cohesion of bacterial genomes. Science 301, 829–832 (2003).

39. Olm, M. R., Brown, C. T., Brooks, B. & Banfield, J. F. dRep: a tool for fast and accurate genomic comparisons that enables improved genome recovery from metagenomes through de-replication. ISME J. 11, 2864–2868 (2017).

40. Ondov, B. D. et al. Mash: fast genome and metagenome distance estimation using MinHash. Genome Biol. 17, 132 (2016).

41. Cavalier-Smith, T., Ema, E. & Chao, Y. Multidomain ribosomal protein trees and the planctobacterial origin of neomura (eukaryotes, archaebacteria). Protoplasma 1–133 (2020).

